# Model-based identification of the crosstalks and feedbacks that determine the doxorubicin response dynamics of the JNK-p38-p53 network

**DOI:** 10.1101/2020.03.10.985994

**Authors:** Laura Tuffery, Melinda Halasz, Dirk Fey

## Abstract

Cellular responses to perturbations and drugs are determined by interconnected networks, rather than linear pathways. Individually, the JNK, p38 and p53 stress and DNA-damage response networks are well understood and regulate critical cell-fate decisions, such as apoptosis, in response to many chemotherapeutical agents, such as doxorubicin. To better understand how interactions between these pathways determine the dynamic behaviour of the entire network, we constructed a data-driven mathematical model. This model contains mechanistic details about the kinase cascades that activate JNK, p38, AKT and p53, and free parameters that describe possible interactions between these pathways. Fitting this model to experimental time-course perturbation data (five time-courses with six time-points under five different conditions), identified specific network interactions that can explain the observed network responses. JNK emerged as an important control node. JNK exhibited a positive feedback loop, was tightly controlled by negative feedback and crosstalk from p38 and AKT, respectively, and was the strongest activator of p53. Compared to static network reconstruction methods, such as modular response analysis, the model-based approach identifies biochemical mechanisms and explains the dynamic control of cell signalling.

## Introduction

Cellular responses to perturbations or drugs are determined by interconnected networks, rather than linear pathways^1^. For example, even within a single pathway such as the ERK pathway, negative feedback can induce a transient response to a sustained stimulus thereby promoting cell proliferation, whereas positive feedback can ensure a sustained response and cell differentiation, even for pulsatile, transient inputs^2^. Further, the response behaviour of one pathway may also be controlled by crosstalks from other pathways. For example, crosstalk from AKT signalling shifts the activation curve of the JNK stress response towards higher tolerable stress inputs, and may even completely disable the JNK bi-stable switch^3,4^.

The JNK, p38 and p53 stress and DNA-damage response networks are of particular interest, because they regulate vital cell fate decisions in response to many chemotherapeutical agents used in cancer therapy^4,5^. Firstly, the stress-activated kinases JNK and p38 can induce programmed cell-death and initiate the cell-cycle arrest by phosphorylating their respective substrates^6–9^. Secondly, the kinase AKT promotes cell-survival and can for example supress the stress response by phosphorylating and inhibiting kinases upstream of JNK and p38^3,10,11^. Thirdly, the transcription factor p53 maintains genomic stability and supresses cancer formation by activating DNA repair proteins, arresting cell growth by pausing the cell cycle, initiating cell-death, and promoting senescence by shortening telomeres^5,7^. Most of these p53 functions require the activation of p53, which is usually achieved by phosphorylation of p53 at sites in its transcriptional domain. Kinases known to target these phosphorylations are amongst others JNK, p38 and AKT. The resulting activation inhibits the p53-MDM2 interaction and MDM2 mediated degradation. In response, the increasing concentration of active p53 induces the expression of p53 target genes, that by driving the mentioned cell fates.

The described p53 activation is induced by many stresses and drugs, including Doxorubicin, which is one of the most frequently used anti-cancer drugs in combination with surgery^12,13^. Doxorubicin is a chemotherapeutic agent from the anthracycline class, acting as a DNA damaging agent. Doxorubicin is part of standard chemotherapy for many cancers, such as breast and ovarian cancer, but is often ineffective. Doxorubicin not only activates p53 but also JNK and p38. But how the dynamic interplay between these pathways controls the Doxorubicin response is not understood.

We hypothesised that Doxorubicin activates JNK, p38 and p53, and that the dynamic behaviour of this response is controlled by specific network interactions between these pathways. To test this hypothesis, we developed a model-based systems identification approach and built a data-driven mathematical model of the JNK, p38, AKT and p53 network. This model contains mechanistic details about the kinase cascades that activate JNK, p38, AKT and p53, and free parameters that describe possible interactions between these pathways. Fitting this model to experimental time-course perturbation data obtained from MCF10A mammary epithelial cells (a cell-line model for non-mutated, non-malignant breast cells) identified the specific network interactions that could explain the observed network responses. JNK signalling activating p53 emerged as a master regulator boosted by positive feedback to amplify the JNK response and tightly controlled by indirect negative feedback and crosstalk from p38 and AKT, respectively.

## Results

### A mathematical model of the JNK, p38, AKT and p53 response dynamics

To investigate the dynamic crosstalks between JNK, p38, AKT and p53 pathways in relation to Doxorubicin treatments, we constructed a mathematical model of ordinary differential equations. A reaction kinetic graph was defined summarising the current knowledge of the network from the literature (Fig 1). In this model, ATM activates two stress-response cascades and p53. Firstly, ATM phosphorylates MKK4 and MKK7, which in turn phosphorylate and activate JNK. Secondly, ATM phosphorylates MKK3 and MKK6, which in turn phosphorylate and activate p38. Other possible intermediates, such as the MAP3Ks upstream of MKK4/7 and MKK3/6 are not included in the model^14^. Further, ATM mediates the phosphorylation, of p53, which happens directly and indirectly^5,15^, but for simplicity, ATM’s effect on p53 is modelled in a single reaction. Both p53 and phospho-p53 induce the expression of MDM2, leading to a negative feedback loop^16^. p53 phosphorylation stabilises p53 by inhibiting MDM2-binding and MDM2-mediated degradation^17^.

**Fig 1:**
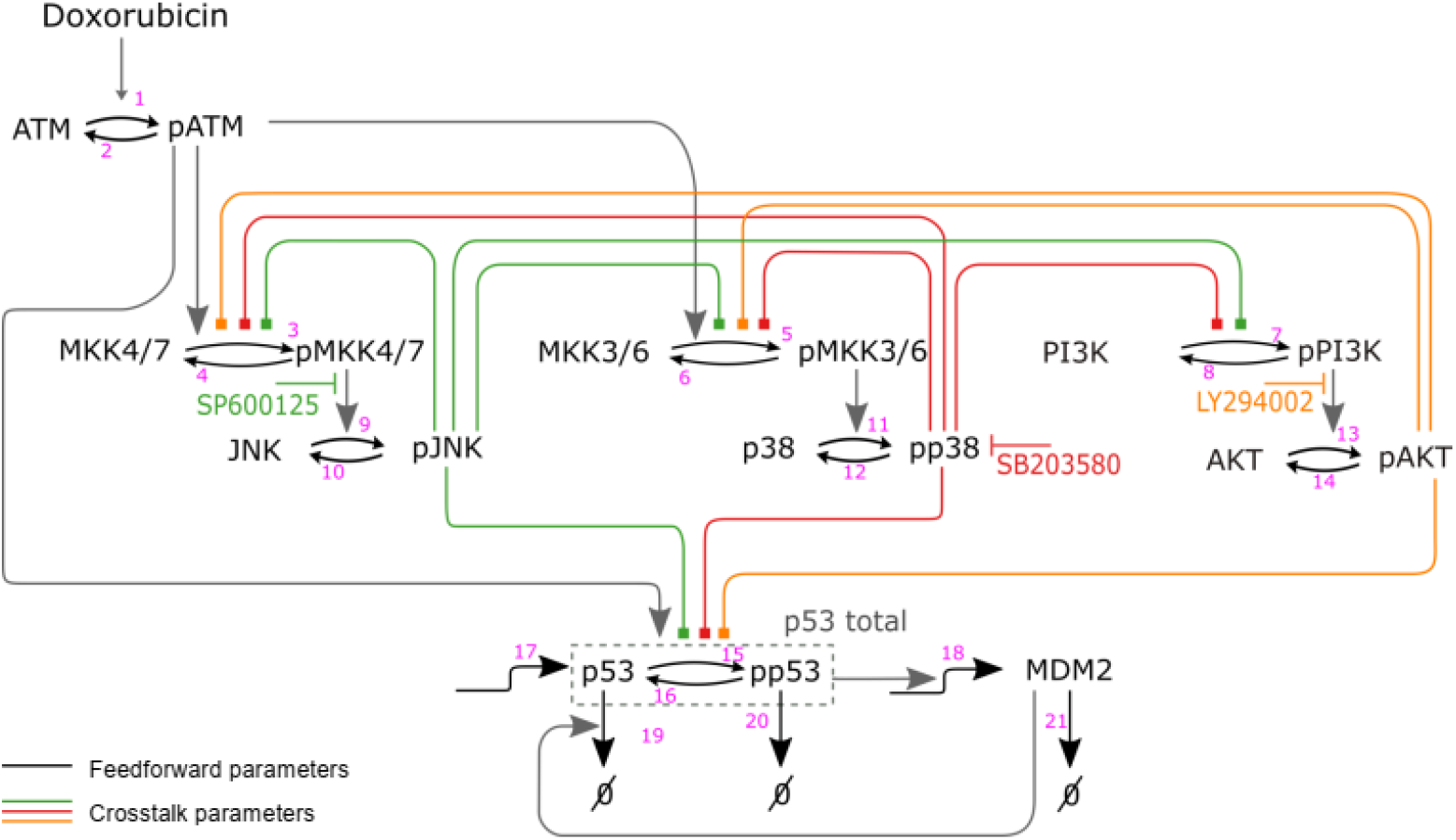
Scheme of the stress and DNA-damage response model. The black arrows indicate biochemical reactions (phosphorylation, dephosphorylation, synthesis and degradation). The grey arrows indicate known canonical pathway interactions from the literature. Coloured lines indicate potential crosstalks (activation or inhibition) from phosphorylated JNK (pJNK), phosphorylated p38 (pp38), and phosphorylated AKT (pAKT) onto the other pathways as indicated in green, red, and orange, accordingly. Numbers on the reactions correspond to the rate laws in Table 1. Red arrows indicate inhibitors for JNK, p38 and PI3K/AKT.

From this reaction kinetic graph a dynamic model in terms of ordinary differential equations (ODEs) can be derived^18^. Table 1 list all the reactions and their rate laws, Table 2 the resulting system of ODEs and their initial conditions.

To limit the complexity of the model and keep the model calibration feasible, the model contains a few simplifying assumptions^4,18^. Firstly, the activation of JNK, p38 and their upstream kinases requires the phosphorylation at two conserved residues. The two phosphorylations were modelled as a single phosphorylation/dephosphorylation cycle. This keeps the model simple and focusses on the active, double-phosphorylated JNK and p38 forms for which we generated experimental data. Secondly, MKK4 and MKK7, and also MKK3 and MKK6, were lumped into two combined model components called MKK4/7 and MKK3/6, respectively. The justification is that the only function in the context of this model is the activation of JNK and p38, respectively. Thirdly, albeit there are many phosphosites on p53, most these phosphorylation sites share a common function, which is the activation and stabilisation of p53^5^. Thus, we only modelled a phosphorylated and an unphosphorylated form of p53.

### Modelling inhibitors

Inhibitors were modelled as additional factors in the activating phosphorylation reactions (Table 1) assuming a non-competitive mechanism^19–21^. For example, the inhibition of JNK is described by the SP term in v9, which is the reaction for phosphorylating and activating JNK. Admissible values for these inhibition factors range from zero (complete inhibition) to one (no inhibition). We adopted the following notation: SP, SB and LY in reactions v9, ap38 and v13, denote the effectiveness of the JNK (SP6001250), p38 (SB203580) and PI3K (LY294002) inhibitors, accordingly. The values of these inhibition factors were estimated from the experimental data as the ratio between the phosphorylated target, for example pJNK, in inhibited versus uninhibited conditions at t=2h (Methods). This yielded the following values SP = 0.27, LY = 0.65, SB = 0.40. The SB203580 inhibitor does not directly inhibit the phosphorylation of p38 but only blocks its catalytic activity. Thus, to verify the effectiveness of the inhibition, we measured the phosphorylation of the p38 substrate MAPKAPK2 (Fig S1), and estimated SB = 0.4.

Because the p38 inhibitor does not directly affect the phosphorylation of p38, the phosphorylation level of p38 does not correspond to p38 activity^22^. In the presence of the inhibitor the phosphorylated p38 is not active. To differentiate phosphorylated form of p38 from the active form, we introduced an additional model variable called ap38 that describes the active form of p38 as *ap*38 = *SB pp*38, where *pp*38 denotes phosphorylated p38.

### Modelling crosstalks

The interactions between the canonical pathways were modelled using crosstalk terms (Fig 1, Table 1). The model assumes that all the crosstalks between the pathways are possible and pJNK, pp38, pAKT mediate these crosstalks. It further assumes that the crosstalks and feedbacks act on the phosphorylation reactions of MKK4/7 upstream of JNK (v3), MKK3/6 upstream of p38 (v5), PI3K upstream of AKT (v7) and p53 (v15). These reactions were chosen for mediating the crosstalks because many known crosstalks act on the level of these MAP2Ks and because these are the first reactions in the model where more upstream crosstalks would take effect in the model^3,4,23^. Mathematically, these crosstalks are described by Hill-type equations that modify the nominal phosphorylation rate laws

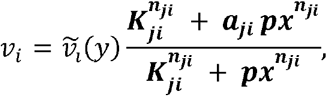

where v_I_ denotes the modified reaction rate taking account of the crosstalks, 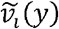 the nominal, unmodified rate law of the phosphorylation reaction, 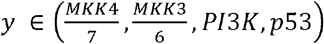 the substrate to be phosphorylated, *px* ∈ (*pJNK*, *pp*38, *pAKT*) the kinase mediating the crosstalk, *K_ji_* > 0 the half-activation constant of the crosstalk, *a_ji_* ∈ [0, ∞) the strength of the crosstalk, *n_ji_* its Hill exponent, and where the indices *j* ∈ (*j*, *p*, *a*) indicate the source of the crosstalk (j for pJNK, p for pp38, a for AKT), and *i* ∈ (3,5,7,15) the number of the reaction they modify (Table 1).

The model allows for crosstalks from each kinase pathway to each other kinase pathway and p53. The aim is to estimate the activation threshold and strength of these crosstalks, whereby *a_ji_* < 1 describe negative crosstalks, *a_ji_* ≈ 1 no crosstalk, and *a_ji_* > 1 positive crosstalks.

### Perturbation experiments generated a rich time-course data set for model calibration

To characterise the network dynamics and calibrate the model, we generated time-course data in response to Doxorubicin +/− inhibitors (Fig S2). MCF10A cells were used as a non-malignant breast cell line model that does not harbour mutations in the modelled pathways. Western blots were used to measure the responses of pJNK, pp38, pAKT and p53 in the presence and absence of the following inhibitors JNK, p38 and PI3K. The data were quantified and normalised as described in the Methods. The resulting dataset comprised at least three biological replicates of five time-courses at six timepoint in four different conditions. This totalled 120 data-points per replicate and build the basis for estimating the parameters.

The perturbation data exhibited the following salient features. Firstly, JNK inhibition strongly inhibited the pJNK response, slightly delayed and reduced the pp38 response, and strongly decreased the p53 response (Fig 2). Secondly, p38 inhibition markedly increased the pJNK and pp38 responses, and slightly reduced the p53 response (Fig 2). That p38 inhibition increased the pp38 response can be explained as follows. The p38 inhibitor SB203580 inhibits p38 catalytic activity by binding to the ATP binding pocket, but does not inhibit phosphorylation of p38 by upstream kinases^22^. The increase of p38 phosphorylation the presence of the inhibitor can therefore be explained by a negative feedback loop and the release of this negative feedback in response to the inhibitor. Thirdly, PI3K inhibition strongly increased the pJNK and pp38 responses and markedly decreased the p53 response (Fig 2). Last but not least, pAKT exhibited very minor changes in response to doxorubicin (Fig 2), albeit pAKT levels were affected by the inhibitors, with the most prominent reductions observed with PI3K and p38 inhibition (Fig 2).

**Fig 2:**
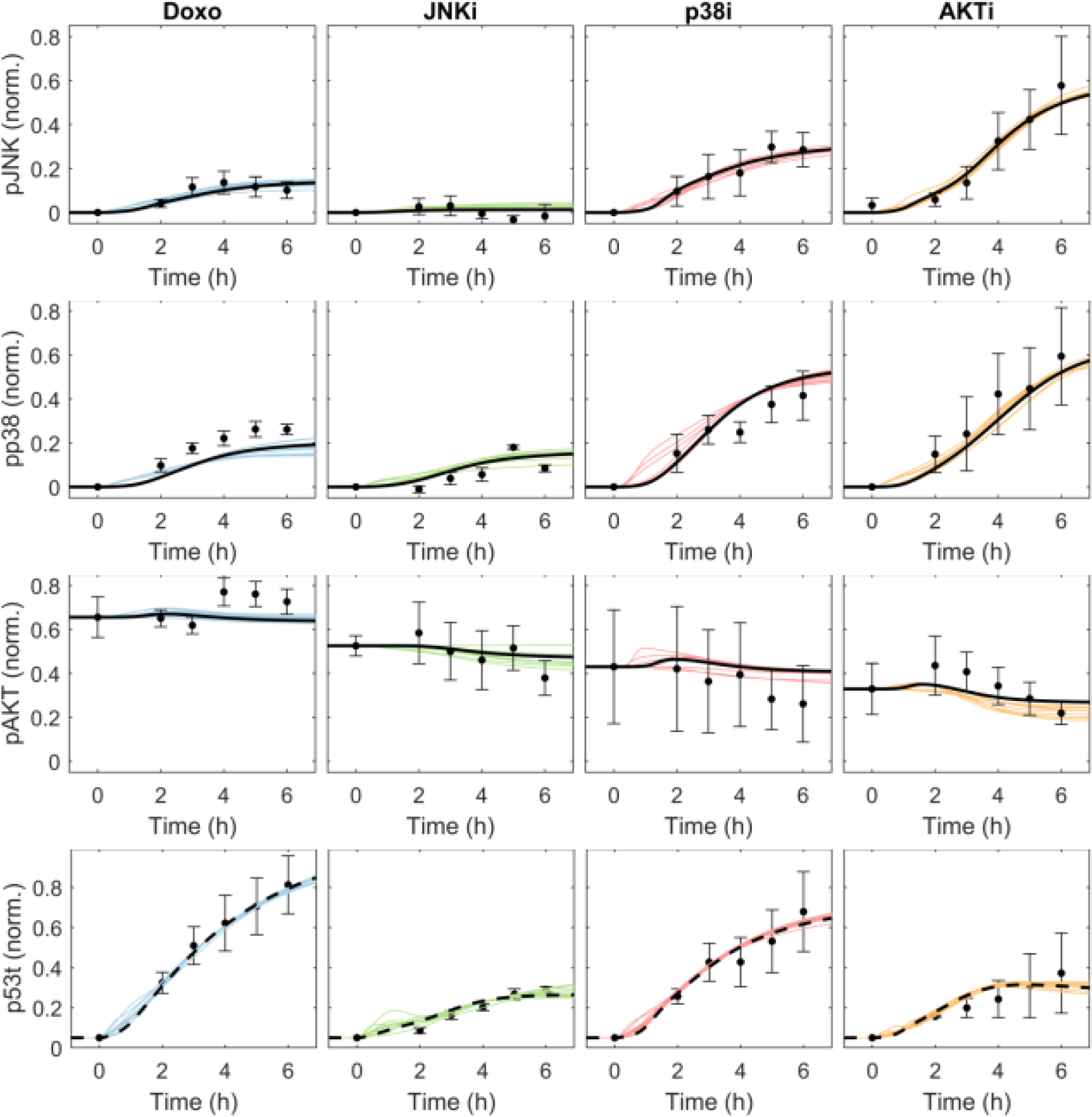
Model fit to the experimental data after the final round of parameter estimation. Lines represent simulations as indicted, circles and error-bars experimental data (mean and standard error of the mean) from three independent experiments. Colours indicate experimental conditions as follows: Doxorubicin treatment alone (control, blue), or in combination with JNK inhibitor (SP600125 at 20μM, red), p38 inhibitor (SB203580 at 100μM, yellow) or PI3K inhibitor (LY294002 at 10nM, purple). Shown are the trajectories over time after Doxorubicin treatment at t=0h for phosphorylated JNK (pJNK), phosphorylated p38 (pp38), phosphorylated AKT (pAKT), and total p53 (p53t).

No direct conclusions should be drawn from these perturbation experiments regarding the network structure^24^. The measured responses constitute so-called global responses that are the consequences of direct and indirect network interactions. For example, the observed impairment of the p53 response in response to the JNK inhibitor could be due to a direct JNK-p53 interaction or an indirect JNK-p38-p53 interaction loop. To detangle direct from indirect effects, formal network reconstruction techniques have to be applied^24–28^. Here, we use the constructed model and parameter estimation to identify the interactions parameters.

### Parameter estimation lead to reliable model simulations fitting the experimental data

The aim of the parameter estimation is to fit the experimental data to the model and estimate all unknown parameters, in particular the crosstalk parameters. To fit the experimental data, we used PEPSSBI with the GSLDC global optimisation algorithm^29^. Biologically feasible parameter bounds were used (*Methods*, Suppl. File 1). Because we had many parameters to estimate, the parameter estimation was performed in a modular fashion divided into four steps. For each step, we performed 100-rounds of independent parameter estimation runs and selected the estimate with the best goodness-of-fit value for the next step. First, we obtained an estimate of the feedforward parameters of the upstream pathways (k1 to k14 and Km1 to Km14) by fitting to the measured pJNK, pp38 and pAKT time-courses in response to Doxorubicin in control conditions without any inhibitors. In this step, the inhibitor data were not used, and the crosstalks were neglected by fixing *a_ji_* = 1. Second, we estimated the crosstalks parameters of the upstream kinase pathways (aj3, aj5, aj7, ap3, ap5, ap7, aa3, aa5, aa7, Kj3, Kj5, Kj7, Kp3, Kp5, Kp7, Ka3, Ka5, Ka7) by fitting the measured pJNK, pp38 and pAKT time-courses for all four conditions. Third, to get a parameter estimate of the downstream MDM2-p53 module (k15, k16, k19, k20, Km15, Km16, Km18 and Km19), we used the best fitting estimates from the previous step, kept these fixed and fitted the measured p53 time-course. Finally, we estimated all the crosstalks parameters (aj3, aj5, aj7, ap3, ap5, ap7, aa3, aa5, aa7, Kj3, Kj5, Kj7, Kp3, Kp5, Kp7, Ka3, Ka5, Ka7, aj15, ap15, aa15, Kj15, Kp15, Ka15) by fitting all the data. Supplementary File 1 contains all parameter estimates.

Figure 2 shows a good correspondence of the model simulations with the experimental data. The simulations from all 100 parameter estimation runs fitted the data well.

### Model based systems identification revealed critical feedback and crosstalk structures

The parameter estimation yielded estimates for the crosstalk parameters of the network. First, we describe the identified crosstalk structure of the best fitting estimate. Then we judge the reliability of the identified structure based on the distribution of all 100 estimated parameter sets.

The best fitting model exhibited a direct positive feedback loop from JNK onto itself, a indirect negative feedback loop in which JNK facilitated the activation of p38, which in turn inhibited the activation of JNK, and negative crosstalk from AKT to JNK (Fig. 3a). All these interactions activated at low concentrations (small *K_ji_* values). p38 activity was controlled by the same indirect negative feedback loop (p38 ⍰ JNK ⍰ p38), a direct positive feedback that only activated at high pp38 concentrations (high value for *K*_*p*5_), and negative crosstalk from AKT activating at low pAKT concentrations. All crosstalks to AKT were very weak (*a*_*j*7_ values close to 1). The p53 response was controlled by strong positive crosstalk from JNK and AKT activating at low pJNK and high pAKT concentrations, respectively, and negative crosstalk from p38 activating at low pp38 concentrations. Figure 3a depicts the inferred network structure of the best fitting model.

**Fig 3:**
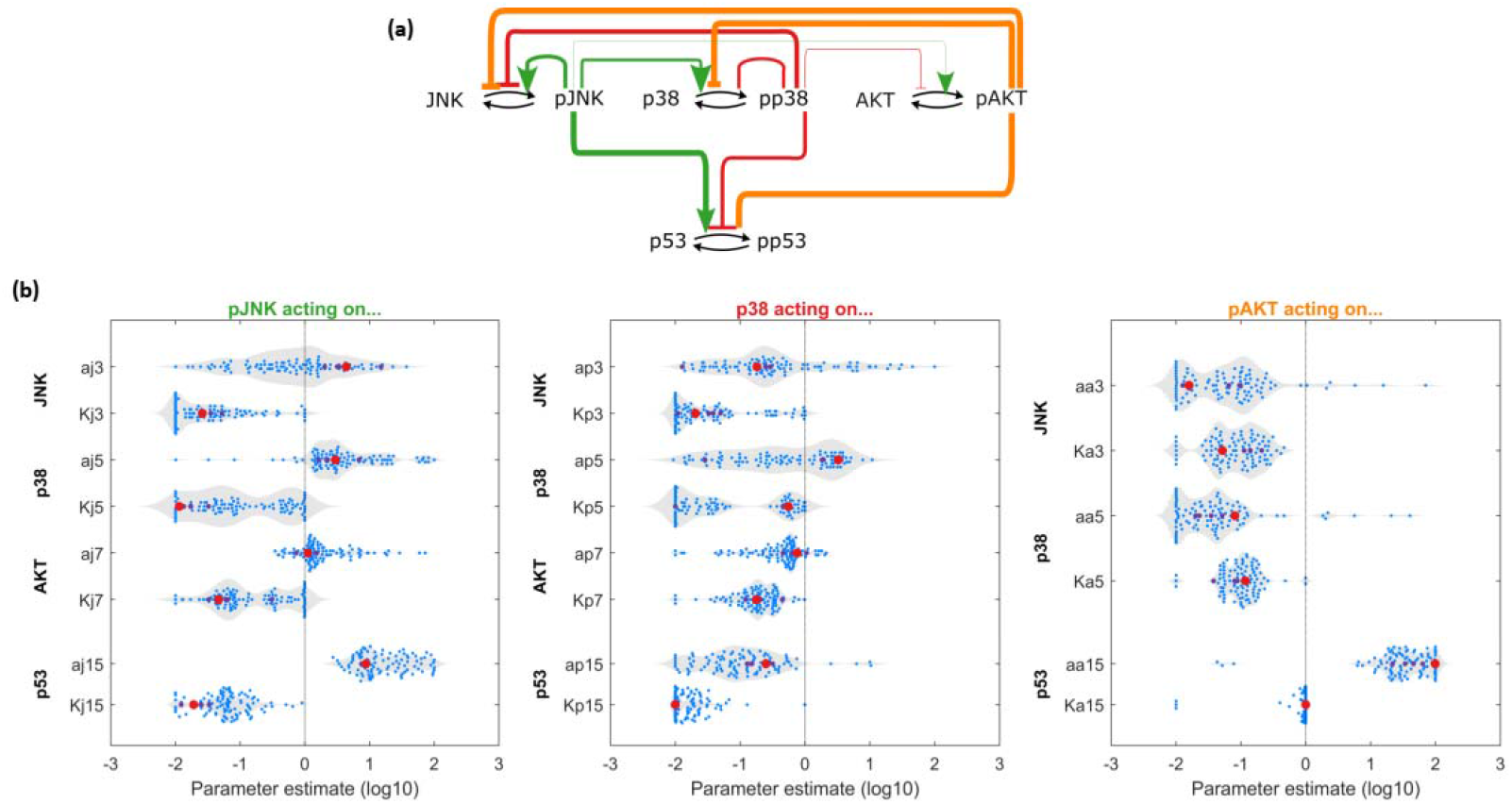
Identified interactions. **(a)** Interaction map of based on the best fitting estimate. Black arrows represent phosphorylation and dephosphorylation of the measured model components. The coloured lines indicate the estimated crosstalks, with the type of the arrowhead indicating the sign of the interaction; ⍰ for activation *log*_10_*a*_*ji*_ > 0, ⍰ for inhibition *log*_10_*a*_*ji*_ < 0. Line thickness is proportional to the strength of the estimated crosstalks |*a_ji_*|. Large arrowheads indicate *K_ji_* < 0.1, small arrowheads *K_ji_* < 0.1. **(b)** Distribution of the crosstalk parameter estimates. Each blue dot represents a parameter estimate from n=100 estimation runs. The red dot indicates the parameter estimate with the best goodness-of-fit value. x-axis is on the log10 scale.

To judge the reliability of the identified network structure we can look at the distribution of the estimated interaction parameters for all 100 estimation runs (Fig. 3b). Some estimates are more dispersed the others. The most consistent estimates are

- Positive crosstalk from JNK to p38 (aj5) that activates at a range of pJNK concentrations (Kj5)
- Negative crosstalk from p38 to JNK (ap3) that activates at low pp38 concentrations (Kp3)
- Negative crosstalk from AKT to JNK (aa3) and p38 (aa5) that activates at intermediate pAKT concentrations (Ka3, Ka5)
- Strong positive crosstalk from JNK to p53 (aj15) that activates at low pJNK concentrations (Kj15)
- Strong positive crosstalk from JNK to p53 (aj15) that activates at low pJNK concentrations (Kj15)Negative crosstalk from p38 to p53 (ap15) that activates at low p38 concentrations (Kp15).
- Strong positive crosstalk from JNK to p53 (aj15) that activates at low pJNK concentrations (Kj15)Strong positive crosstalk from AKT to p53 (aa15) that activates at high pAKT concentrations (Ka15).

The estimated crosstalk strengths from p38 and JNK to AKT (aj7, ap7) were concentrated near zero on the log scale indicating that these crosstalks are very weak.

The most dispersed estimates were the direct JNK feedback (aj3), the direct p38 feedback (ap5), which depending on the parameter estimation run, could take both positive and negative values. Therefore, we analysed these estimates in more detail.

### Specific correlations within the estimated crosstalk parameters revealed mutually exclusive crosstalk patterns between JNK and p38

The large dispersion of the direct JNK and p38 feedback parameters (aj3 and ap5) indicates the practical non-identifiability of these parameters^30^. In situations where unique parameter values cannot be identified, they are often correlated with other parameters. When such correlations exist, relations between different parameters could still be identifiable^30^. For example, the identified value of the JNK feedback strength (aj3) could depend on the crosstalk strength from p38 (ap3) as both crosstalks act on the same reaction (v3). To confirm this hypothesis, we performed a correlation analysis and looked at the relationships between the estimated parameter values.

To better understand the results of the parameter estimation, we investigated the relationships between the parameters in a correlation analysis. We observed a strong negative correlation between the direct JNK feedback parameter aj3 and the p38-to-JNK crosstalk parameter ap3 (Fig 4a). This anti-correlation is consistent with models that feature either a strong JNK feedback or a strong crosstalk from p38, but not both. Indeed, a scatter plot of the estimates for aj3 versus ap3 (Fig 4b) confirms that a unique value for each parameter was not practically identifiable, but the relationship between these two parameters was. The scatter plot shows a mutually exclusive pattern where JNK features either a strong positive crosstalk from p38 (top-left corner) or positive feedback onto itself (bottom right corner).

**Fig 4:**
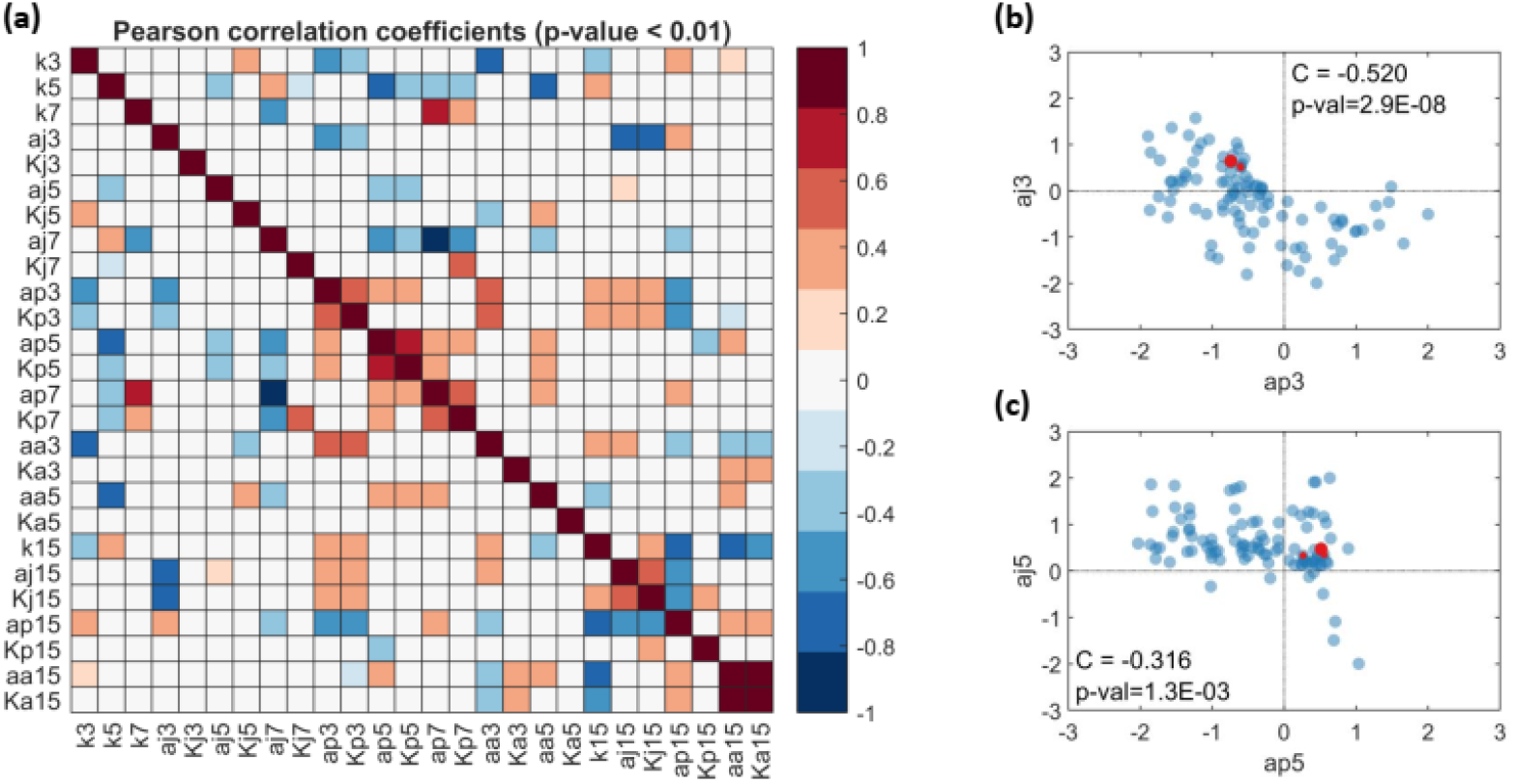
Parameter correlations based on n=100 estimates. (a) Heatmap of the Pearson correlation coefficients for parameter estimates of the crosstalk parameters from the last round of parameter estimation. (b) Scatterplot of *a*_*j*3_ versus *a*_*p*3_, and (c) *a*_*j*5_ versus *a*_*p*5_. C denotes the Pearson correlation coefficient.

A similar, albeit weaker, pattern was observed between the direct p38 feedback ap5 and the JNK-to-p38 crosstalk aj5 (Fig 4c). Most parameter estimates featured a negative p38 feedback accompanied by a positive JNK-to-p38 crosstalk (top right), whereas negative JNK-to-p38 crosstalk was predominantly accompanied by positive p38 feedback.

### Modular Response Analysis identified a similar network structure

To benchmark the inferred network, we compared our model-based results to a well-established network inference method Modular Response Analysis (MRA). The strength of MRA is that it can identify the exact network structure between all measured nodes given that each node has been perturbed 1-by-1 in experiments and measured. Because not all measured nodes were perturbed in our network (p53 was not perturbed), we applied a recent extension of MRA (Bayesian MRA) that can deal with incomplete perturbation data. Unlike our model-based systems identification, MRA cannot identify the reaction-kinetic mechanistic details of the crosstalks. Nevertheless, MRA allows us to reconstruct the connectivity of the network and compare the result to our model-based inference.

We applied Bayesian MRA (BMRA) on the measured nodes (see Methods for details) and compared the results to the estimated crosstalk parameters *a_ji_*, Figure 5. BMRA uses the so-called global response coefficients, that is the measured relative changes of the stead-state responses to a set of perturbations, to infer the connectivity of the network in terms of so-called local response coefficient *r_ji_*. A non-zero *r_ji_* means node j is directly regulated node j. For technical reasons, BMRA cannot identify direct feedbacks, because the diagonal elements of the local response coefficients have to be normalised *r_ii_* = 1. Therefore, only off-diagonal interactions (crosstalks) can be compared.

**Fig 5:**
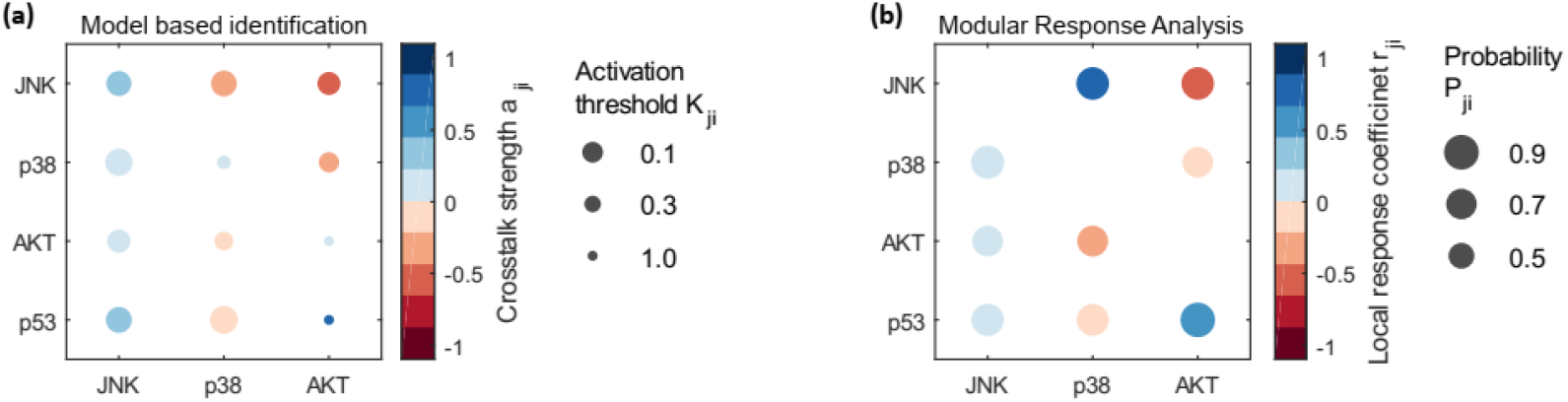
Comparison of the model-based systems identification result with Bayesian Modular response analysis. Each circle represents a crosstalk interaction from the node on the x-axis to the node on the y-axis. (a) Model-based result. The color of the circle represent the sign and strength of the interaction *a_ji_* as indicated in the colorbar legend, the size the activation threshold *K_ji_*. (b) BMRA result. The color represent the sign and size of the estimated local response coefficient *r_ji_*, the size the associated estimated probability of a nonzero interaction.

The BMRA results were largely concordant with the model-based results; 8 out of 9 crosstalk interactions were estimated to have the same sign for both methods (Fig. 5). For both methods, p38 was positively regulated by JNK and negatively by AKT, p53 was positively regulated by JNK and AKT and negatively by p38, AKT was positively regulated by JNK and negatively regulated by p38. The only discrepancy was observed for the p38-to-JNK crosstalk. JNK was negatively regulated by AKT for both methods, but in contrast to the model which inferred a positive regulation by p38, BMRA inferred a negative regulation. This discrepancy for the p38-to-JNK crosstalk might be explained by the limitations of BMRA, which assumes small perturbations, is sensitive to measurement noise, and the requires that the responses are measured in steady state. In contrast, our model can simulate the dynamic time-courses and matches these experimental data well (Fig. 2). The experimental data show a strong increase of the dynamic pJNK response under p38 inhibited conditions, which is indicative of a negative crosstalk. Accordingly, our model-based approach identified a negative crosstalk from p38 to JNK (a_p3<0).

### Structural sensitivity analysis established JNK as a master regulator of the network response dynamics

Next, we wanted to better understand how the crosstalks and feedbacks control the dynamic behaviour of system. Therefore, we performed a structural sensitivity analysis in which each crosstalk and feedback was removed one-by-one, and the such perturbed model was simulated. In contrast to inhibitor experiments, which block all outgoing interactions from the inhibited node, this simulation-based approach reveals the control of each individual crosstalk.

Figure 6a shows the simulated responses for each perturbation. Overall, the largest sensitivities were observed for interactions relating to JNK (Fig 6b). Five of the seven largest sensitivities were related to JNK (aj3, positive feedback; aj5, ap3; negative feedback via p38; aa3, negative crosstalk from AKT; aj15, JNK activating p53). The remaining two large sensitivities were related to AKT (a_a5, negative crosstalk to p38; a_j15, positive crosstalk to p53). These can be interpreted as follows. The JNK positive feedback amplifies the pJNK response, and also had the greatest impact on p53 activation compared to all other upstream crosstalk parameters. The JNK negative feedback loop via p38 (aj5>1, ap3<0) serves to attenuate the JNK response while concomitantly amplifying the p38 response. AKT was not notably controlled by any crosstalk parameter, while individual AKT-to-JNK and AKT-to-p38 crosstalks served to attenuate the JNK and p38 responses, respectively. In summary, large sensitivities were observed for crosstalk parameters regulating JNK or mediating JNK-dependent effects.

**Fig 6:**
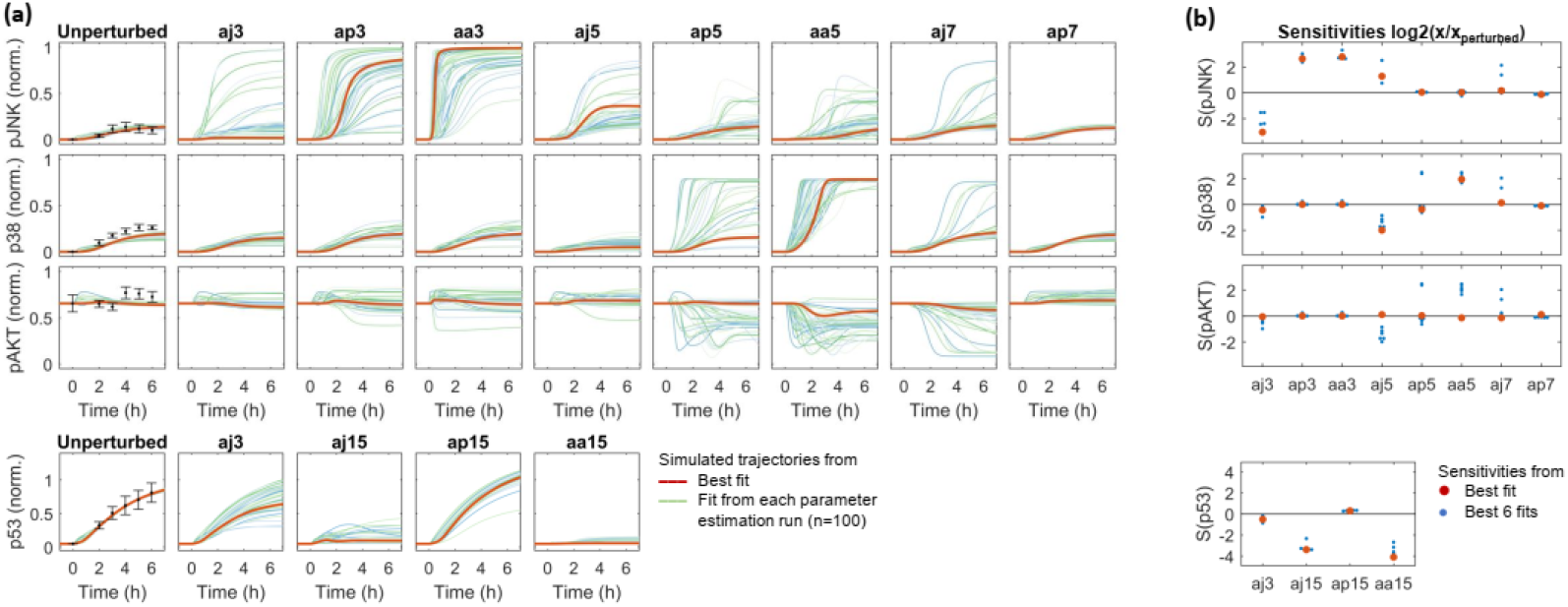
Structural sensitivity analysis of the model. Each interaction in the model was removed one by one and the model was simulated for all parameter estimates. (a) The simulated trajectories for the unperturbed and perturbed models as indicated for all 100 parameter estimates. For the unperturbed model, the experimental data are shown with dot and errorbars representing mean and standard error. (b) Calculated sensitivity coefficients for a subset of the perturbations as indicated for the six best fitting parameter estimates. Red dots represent the best fitting model.

## Discussion

We have shown that Doxorubicin activates the JNK, p38 and p53 network in MCF10A cells, and constructed a data-driven mathematical model of these network dynamics. This model describes the mechanism of the network interactions, and contained free parameters describing possible interactions between the modelled pathways. Estimating these parameters explained the experimentally measured time-course perturbation data and identified the specific crosstalk patterns that control the Doxorubicin response. The identified model shows that that direct JNK positive feedback serves to amplify the JNK doxorubicin response but is tightly controlled by indirect negative feedback via p38 and negative crosstalk from AKT. The role of the indirect negative feedback is to attenuate the JNK response while concomitantly amplifying the p38 response. JNK, not p38, was a major activator of p53, which underlines the importance of controlling the JNK response. AKT also contributed positively to p53 activation. In the following we discuss the relevance of each crosstalk.

The most consistent model estimates were positive crosstalk from JNK to p38 and negative crosstalk from p38 to JNK constituting the indirect negative feedback loop, negative crosstalks from AKT to both JNK and p38, and positive crosstalks from JNK and AKT to p53.

The negative crosstalks from p38 and AKT to JNK are well supported by the literature and their biochemical mechanisms have been established. Chemical inhibition or knock-out of p38 caused JNK hyperactivation in several contexts, demonstrating that p38 negatively regulates JNK ^31–36^. Mechanisms concern both posttranslational and transcriptional processes depending on stimulus and include the decreased phosphorylation of MKK4^33^ and MKK7 and the upregulation of the JNK dual specificity phosphatase DUSP1^36^. In single cells, p38 cross inhibition supresses JNK activation dynamics to ensure the right amount of fractional cell killing^37^. The inhibition of JNK by AKT is also well established. AKT can phosphorylate and inhibit the JNK upstream kinases MKK4 and MKK7 at S80^23^ and T385^3^, respectively.

The activation of p53 by JNK and AKT is also well supported by the literature. JNK can phosphorylate and stabilise p53 at S20^38^ and T81^39,40^, and doxorubicin induced JNK mediated p53 activation and cell death in SY5Y cells^3^. Although p53 can also be phosphorylated by p38 at S33^41–43^ and S46^41^, our p38 inhibition had only minor effects on the p53 response. Accordingly, the identified p38-p53 crosstalk was weak in our model. In contrast, the identified AKT-p53 crosstalk was strong. Literature suggests that this positive crosstalk from AKT to p53 is likely not direct; p53 stabilization in response to DNA damage depends on AKT mediated phosphorylation and inactivation of GSK3 that normally phosphorylates and inactivates MDM2^44^. Thus, AKT activity inactivates GSK3 and MDM2 thereby preventing p53 protein degradation and facilitating p53 accumulation. Summarizing, these observations suggest that a direct p38-p53 crosstalk plays a minor role in the doxorubicin response, and that p53 activation is mainly controlled by crosstalk from JNK and AKT.

The identified positive crosstalk from JNK to p38 positive is novel, and no potential mechanisms have been reported. However, JNK can exhibit a positive feedback by phosphorylating MKK7 at T66 and T83, thereby enhancing its own activation^3^. Considering that the p38 upstream kinases MKK3 and MKK6 are structurally very similar to MKK7, is conceivable that JNK might phosphorylate p38 upstream kinases MKK3 and MKK6, thereby enhancing the activation of these kinases. Another possible mechanism for mediating crosstalks are dual specificity phosphatases (DUSPs) that are often induced by JNK and/or p38 signalling and that can act as negative and positive regulators^4^. An example for the latter is DUSP22 which has been shown to enhance JNK activation^45^.

In contrast to the network interactions discussed above, the estimates of the JNK feedback were more variable and could take positive or negative values. However, all six best fitting model exhibited positive feedback with low half activation constants. In contrast, estimates that featured a negative feedback strength exhibited large values for the half activation, thus diminishing the negative effect. Further, JNK positive feedback under stress conditions is consistent with the literature. JNK positive feedback was observed under many stress conditions and cell types^4,14,46–50^. Concerning the mechanism, experiments in SY5Y cells have shown that JNK phosphorylates it upstream activator MKK7 at the T66 site (that flanks the D domain responsible for substrate docking), which increased JNK phosphorylation at the catalytic site (probably by relieving a N-terminal auto-inhibitory domain). These literature data corroborate the results of best fitting models and the identification a JNK positive feedback.

Controlling the JNK activation dynamics seems particularly important considering that JNK activation can exhibit two phases; an early, transient phase that promotes cell survival and a late, sustained phase that promotes apoptosis^4,51,52^. In response to cell stress, such as doxorubicin, the proapoptotic activation of JNK represent a critical signalling mode. The emerging picture is that proapoptotic JNK signalling is controlled by many loops. Once activated, the JNK positive feedback promotes apoptosis. It makes sense that this positive loop must be tightly controlled and kept in check by negative feedback loops and crosstalks. In this context the function of p38 and AKT signalling is attenuate the JNK response and control the levels of cell death^3,37^. For example, in singe cells, cross-inhibition of JNK by p38 lead to right amount of fractional cell killing, while blockade of p38 signalling lead to hyperactivation of JNK in all cells and eradication of the entire cell population^37^.

The developed model-based approach for systems identification has several advantages over Modular Response Analysis and similar approaches^24–26,53–55^. In contrast to standard network inference method, the model-based approach describes the biochemical mechanisms of the identified network interactions and can be used to analyse the dynamics of the system and its control. Applying BMRA to our data yielded very similar results in terms of the identified network structure. However, BMRA infers these interactions from steady-state data, assumes small perturbations, does not yield a mechanistic dynamic model, and can thus not explain the system’s activation dynamics. In contrast out model-based approach focusses on gaining mechanistic insight and explaining the system dynamics. That the shape of the JNK response dynamics is important was observed previously, where the steepness of the pJNK dose-response was prognostic of patient survival^3^. Similar observations were made for the p53 activation dynamics^1,5,56–58^. Rapid activation of p53 leading to higher p53 amplitudes promotes cell death^57^. In those time-dependent, highly dynamic situations a model-based approach is more appropriate.

The model also suggests optimal designs for follow up experiments. The estimates of the JNK feedback were particularly varied, which could be resolved by measuring the upstream kinases MKK4 and MKK7 to estimate the exact feedback strength. The JNK and p38 crosstalks could activate at either low or high pJNK and pp38 concentrations, which could be resolved by recording a dose response to precisely estimate the half-activation constants. These model-based experimental designs could be used in follow up-experiments to remove some of the observed parameter uncertainties. The model could also be used to identify and explain crosstalks for other drugs and cell types. An interesting open question is how context specific theses crosstalks are and whether they change depending on cell type and drug stimulus^59,60^. Generally, the developed model-based systems identification approach could also be applied to other models and crosstalk systems. Considering that a dynamic model pJNK responses was prognostic of patient survival^3^, it would also be interesting to test whether our model can predict patient-specific response to doxorubicin containing chemotherapy, for example in breast cancer. An interesting question is whether the pJNK response alone or in combination with p38 and p53, is predictive of clinical drug-responses and/or prognostic of relapse and patient survival.

## Supporting information

Main and Supplementary Figures

Table model equations

Parameter estimation results

## Acknowledgements

This project has received funding from the European Union’s Horizon 2020 research and innovation programme under grant agreement No 754923 and Irish Research Council under grant agreement No GOIPG/2015/3850. The material presented and views expressed here are the responsibility of the authors only. The EU Commission takes no responsibility for any use made of the information set out.

## Author Contributions

Conceptualisation: LT and DF; Methodology: LT, MH and DF; Investigation: LT and DF; Formal Analysis: LT and DF; Data curation: DF; Resources: MH and DF; Writing – original draft; LT; Writing – review and editing: MH and DF; Funding Acquisition: LT and DF; Supervision: MH and DF.

## STAR Methods

### Lead Contact and Materials Availability

Further information and requests for resources and reagents should be directed to and will be fulfilled by the Lead Contact, Dirk Fey. This study did not generate any new unique reagents.

### Experimental Model and subject details

#### Cell culture

The MCF10A non-tumorigenic mammary epithelial cell line was cultured in DMEM F-12 (Gibco, Cat. No. 11330032) supplemented with 5% heat inactivated horse serum (Gibco, Thermo Fisher Scientific, Cat. No. 16050122), 20μg/L of EGF (PeproTech, Cat. No. 100-47), 500ng/L of hydrocortisone (Sigma-Aldrich, Cat. No. H4001), 100μg of cholera toxin (Sigma-Aldrich, Cat. No. C8052 SIGMA) and 10mg/mL of insulin (Sigma-Aldrich, Cat. No. 91077C). The inhibitors used were: JNK inhibitor (20μM SP00125; SelleckChem, Cat. No. S1460), p38 inhibitor (10nM SB203580; Calbiochem, Cat. No. S1105) and PI3K inhibitor (10μM LY294002; SelleckChem, Cat. No. S2929).

### Method Details

#### Western blotting

Western blots were prepared from cell lysates after using the following lysis buffer: 20mM tris-HCl (pH 7.5), 150 mM NaCl, and 1% Triton X-100 and 1mM MgCl2. The lysis buffer was complemented with cOmplete Mini Protease Inhibitor Cocktail (Roche, Cat. No. 11 836153 001) and PhosSTOP Phosphatase Inhibitor Cocktail (Roche, Cat. No. 4906837001). SDS-polyacrylamide gel electrophoresis (SDS-PAGE) and Western blotting were performed using 10% acrylamide gels with the Mini-PROTEAN® Tetra Cell Systems from Bio-Rad. SuperSignal Femto Chemiluminescent Substrate (Thermo Scientific, Cat. No. 34095) was used to develop Western blots, which were imaged using the Advanced Molecular Vision Chemiluminescence Imaging System. Quantitative Western blotting was performed using multi-strip Western blotting^61^. The antibodies used for Western blotting were from Cell Signalling Technology: p-JNK (Thr183/Tyr185, Cat. No. 9251), total JNK (Cat. No. 9252), p-P38 (Thr180/Tyr182, Cat. No. 9211), total p38 (Cat. No. 9212), total p53 (Cat. No. 9282), p-AKT (Ser473, Cat. No. 9271), total AKT (Cat. No. 9272), p-MAPKAPK2 (Thr334, Cat. No. 3041), GAPDH (Cat. No. 2118).

### Quantification and statistical analysis

#### Quantification and normalisation of time-course data

Western blots were quantified using the ImageJ software (version 1.51j8)^62^. Three pAKT datapoints diverged substantially from the others and have been excluded as outliers and have not been quantified. These are: Replicate 3, 3h control (Fig S2C); Replicate 4, 4h control and 4h JNK inhibitor (Fig S2D). Data normalisation: First, the quantified Western blot data were normalised by dividing the measured values with the GAPDH loading control. Second, to have comparable values between the different inhibitor experiments the phosphorylation time courses were aligned using sum-normalisation as explained previously^63^. The reference time-course for this sum-normalisation was the measured Doxorubicin time course in the absence of inhibitors (hereafter referred to as Doxorubicin control), which was recorded in all experiments and therefore a suitable reference. Consequently, for each experiment, all recoded values were divided by the sum of the Doxorubicin control time course^63^. We also assumed that the phosphorylation levels before Doxorubicin treatment are negligible and subtracted the value of the zero time point from all the other values of the measured time course. Next, to obtain values between zero and one for each species, we normalised by dividing with the maximum value of the species over all the experiments and conditions. These zero-one normalised datasets have been used for the parameter estimation of the model. The complete set of numerical values for these data can be found in the PEPSSBI file (Suppl. File 1)

#### Calculating the steady state initial conditions

The steady-state initial condition prior to Doxorubicin treatment was calculated as follows. From *doxo* = 0 follows that all phosphorylated protein concentrations are zero and therewith that all phosphorylation and dephosphorylation rates are zero. In particular, with *v*16 = *v*17 = 0 and *pp*53 = 0 the ODEs for p53 and MDM2 become

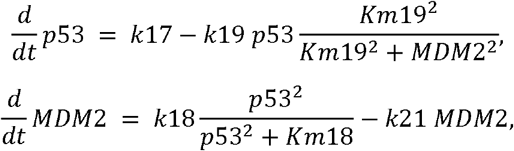

see Table 1. Solving for the steady-state 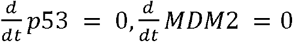 yields

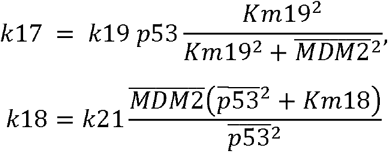

where we assume 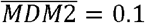 for the MDM2 steady-state and 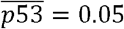 for the p53 steady-state based on our measured data for *doxo* = 0.

Similarly, we calculated the non-zero PI3K and AKT steady states by setting *v*7 − *v*8 = 0 and *v*13 − *v*14 = 0 and using the conserved moieties for the phosphorylation levels (*AKT* + *pAKT* = 1, *PI3K* + *pPI3K* = 1), yielding

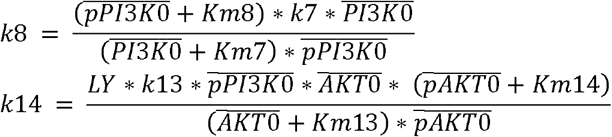

with

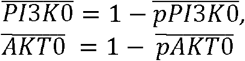

where 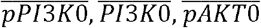 and 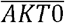 denote the initial steady-state values of pPI3K, PI3K, pAKT and AKT, correspondingly. All other parameters as defined in Table 1. 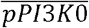 was set to 0.5 and 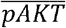 to 0.4, 0.4, 0.28 and 0.23 for the control, JNK inhibited, p38 inhibited and AKT inhibited conditions, correspondingly, according to our measured pAKT data.

#### Parameter estimation and parameter bounds

Parameters were estimated using the *Parameter Estimation Pipeline for Systems and Synthetic Biology* (PEPSSBI) minimising the sum-of-squares objective function with the GSLDC global optimisation algorithm^29^. Lower and upper parameter bounds were specified based on biologically reasonable ranges and information from the literature as follows. For phosphorylation reactions we estimated the rate constant in the range *k* ∈ [0.001,10], and for dephosphorylation in the range *k* ∈ [0.0001,10000]. All Km’s were estimated in the range *Km* ∈ [0.1,2]. The MDM2 half-life is about 30 minutes^64^. Using the half-life equation 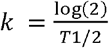, where T1/2 denotes the half-life and k the degradation rate constant, we fixed the MDM2 degradation rate constant at *k*21 = 1.38 1/*h*. The p53 half-life in unstimulated, unphosphorylated conditions is about 5 minutes and increases several-fold when stimulated and phosphorylated^65^. Accordingly we allowed for very rapid MDM2 mediated degradation of p53 within the parameter bound *k*19 ∈ [8, 40], followed by a 10 to 100 fold decrease in degradation rate upon p53 phosphorylation using the parameter bound *kdeg2* ∈ [0.01, 0.1]. For the bounds of the crosstalks parameters we allowed positive or negative effects up to a 100-fold change *a_ji_* ∈ [0.01,100]. The complete parameter estimation setup including experimental data, data normalisation, parameters and parameter bounds, equations and fitting options are specified in the PEPSSBI file (Suppl. File 1).

#### Bayesian modular response analysis

The values of the local response coefficients representing the high-level network structure and their probabilities were estimated using BMRA [REF]. Global response coefficients were calculated according to the

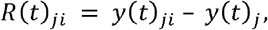

where *R_ji_* denotes the global response coefficient of node *j* ∈ {*pJNK*, *pp38*, *pAKT*, *p53*} to perturbation *i* ∈ {*JNK*, *p38*, *AKT*} (doxorubicin + inhibition of JNK, p38, or AKT), *y_ij_* the measured response of node *j* to perturbation *i*, *y_j_* the measured response of node *j* in control conditions (doxorubicin), and *t* ∈ {5*h*, 6*h*} the timepoint used. Because several replicate measurements were available, the mean was used 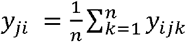, which improves the accuracy of the MRA estimates [REF]. Further, it was assumed that both the 5h and 6h time-points were steady state. Both timepoints were used as observations for BRMA, yielding a 4×6 global response matrix [*R*(5*h*)_*ji*_ *R*(6*h*)_*ji*_], and increasing the precision of the BMRA estimates. The following hyper-parameters were used number of Gibbs samples: noit=10,000, burn-in: burnin=5,000, number of repeats: times=10. The code is provided in Suppl. File 1.

#### Structural sensitivity analysis

Structural sensitivity analysis was performed in Matlab. Each crosstalk was removed one at a time by setting log *a_ji_* = 0. The such perturbed system was simulated and compared to the unperturbed simulation. The sensitivity coefficients were calculated at *t* = 10*h* approximating steady-state using

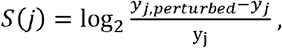

where *s*(*j*) denotes the sensitivity of node *j* ∈ {*pJNK*, *pp38*, *pAKT*, *p53*}, *y_j,perturbed_* the value of the simulated perturbed trajectory of node *j* at timepoint *t* = 10*h*, and *y_j_* the value of the simulated unperturbed trajectory of node *j* at timepoint *t* = 10*h*. The code is available in Suppl. File 1.

### Data and code availability

The authors declare that the data supporting the findings of this study (including mathematical models, experimental data, and parameter estimates) are available within the paper and its supplementary information files (model code: PEPSSBI files, raw images: png files; quantification: Excel table; parameters: Excel table). In addition, the calibrated best-fitting model will be available from the EMBL-EBI BioModels repository (http://www.ebi.ac.uk/biomodels/) under accession number [to be obtained].

---

**Supplementary figure S1: Western blot of phospho-MAPKAPK2 to verify the effect of p38 inhibitor.** MCF10A have been treated with 1μM Doxorubicin after a 30 min pre-treatment of SB203580 at 100μM. Shown are two of the 6 replicates used for calculating the SB203580 inhibition factor.

**Supplementary figure S2(a): Western blotting of experiment number 1.** MCF10A cells were plated at 75.000 cell/mL, pre-treated with 20μM JNK inhibitor (SP600125) for 30 minutes and treated with 1μM Doxorubicin for 0-0.25-0.5-0.75-1-2-3-4-5-6-7-8 hours. The Western blots were performed on the whole cell lysate using the following antibodies: p-JNK (Thr183/Tyr185), total JNK, p-p38 (Thr180/Thy182), total p38, total p53 and GADPH.

**Supplementary figure S2(b): Western blotting of experiment number 2.** MCF10A cells were plated at 75.000 cell/mL, pre-treated with 20μM JNK inhibitor (SP600125), or 100μM p38 inhibitor (SB23580) for 30 minutes, and treated with 1μM Doxorubicin for 0-2-3-4-5-6 hours. The Western blots were performed on the whole cell lysate using the following antibodies: p-JNK (Thr183/Tyr185), total JNK, p-p38 (Thr180/Thy182), total p38, total p53, p-AKT (Ser473), total AKT and GADPH.

**Supplementary figure S2(c): Western blotting of experiment number 3.** MCF10A cells were plated at 75.000 cell/mL, pre-treated with 20μM JNK inhibitor (SP600125), or 100μM p38 inhibitor (SB23580) for 30 minutes, and treated with 1μM Doxorubicin for 0-2-3-4-5-6 hours. The Western blots were performed on the whole cell lysate using the following antibodies: p-JNK (Thr183/Tyr185), total JNK, p-p38 (Thr180/Thy182), total p38, total p53, p-AKT (Ser473), total AKT and GADPH.

**Supplementary figure S2(d): Western blotting of experiment number 4.** MCF10A cells were plated at 75.000 cell/mL, pre-treated with 20μM JNK inhibitor (SP600125), or 100μM p38 inhibitor (SB23580) for 30 minutes, and treated with 1μM Doxorubicin for 0-2-3-4-5-6 hours. The Western blots were performed on the whole cell lysate using the following antibodies: p-JNK (Thr183/Tyr185), total JNK, p-p38 (Thr180/Thy182), total p38, total p53, p-AKT (Ser473), total AKT and GADPH.

**Supplementary figure S2(e): Western blotting of experiment number 5.** MCF10A cells were plated at 75.000 cell/mL, pre-treated with 10nM PI3K inhibitor (LY294002) for 30 minutes, and treated with 1μM Doxorubicin for 0-2-3-4-5-6 hours. The Western blots were performed on the whole cell lysate using the following antibodies: p-JNK (Thr183/Tyr185), total JNK, p-p38 (Thr180/Thy182), total p38, total p53, p-AKT (Ser473), total AKT and GADPH.

**Supplementary figure S2(f): Western blotting of experiment number 6.** MCF10A cells were plated at 75.000 cell/mL, pre-treated with 10nM PI3K inhibitor (LY294002) for 30 minutes, and treated with 1μM Doxorubicin for 0-2-3-4-5-6 hours. The Western blots were performed on the whole cell lysate using the following antibodies: p-JNK (Thr183/Tyr185), total JNK, p-p38 (Thr180/Thy182), total p38, total p53, p-AKT (Ser473), total AKT and GADPH.

**Supplementary figure S2(g): Western blotting of experiment number 7.** MCF10A cells were plated at 75.000 cell/mL, pre-treated with 100μM p38 inhibitor (SB23580), or 10nM PI3K inhibitor (LY294002) for 30 minutes, and treated with 1μM Doxorubicin for 0-2-3-4-5-6 hours. The Western blots were performed on the whole cell lysate using the following antibodies: p-JNK (Thr183/Tyr185), total JNK, p-p38 (Thr180/Thy182), total p38, total p53, p-AKT (Ser473), total AKT and GADPH.

**Supplementary figure S3: Identified interaction maps for the best fitting models 2-5** (parameter estimates in data rows 2-5, Excel file: “Parameter bounds and estimation results.xlsx”, Sheet: “Run 4”). **(a)** 2^nd^ best fitting parameter estimate; **(b)** 3^rd^ best fitting parameter estimate; **(c)** 4^th^ best fitting parameter estimate; **(d)** 5^th^ best fitting parameter estimate. Black arrows represent phosphorylation and dephosphorylation of the measured model components. The coloured lines indicate the estimated crosstalks, with the type of the arrowhead indicating the sign of the interaction; ⍰ for activation *log*_10_*a*_*ji*_ > 0, ⍰ for inhibition *log*_10_*a*_*ji*_ < 0. Line thickness is proportional to the strength of the estimated crosstalks |*a_ji_* |. Large arrowheads indicate *K_ji_* < 0.1, small arrowheads *K_ji_* > 0.1.

**Supplementary figure S4: Distribution of the parameter estimates for each round of parameter estimation: (a)** Round 1; **(b)** Round 2; **(c)** Round 3; **(d)** Round 4. Each blue dot represents a parameter estimate from n=100 estimation runs. The red dot indicates the parameter estimate with the best goodness-of-fit value. x-axis is on the log10 scale.

**Supplementary figure S5: Distribution of the sensitivities for all n=100 estimates from the last round of parameter estimation.** Each interaction in the model was removed one by one and the model was simulated for all parameter estimates (see Fig. 6) and the sensitivity coefficients were calculated. The sensitivities were calculated for all parameter estimates, i.e. each data row in Sheet: “Run 4”, Excel file: “Parameter bounds and estimation results.xlsx”. Each blue dot represents the sensitivity of a parameter estimate (n=100 estimation results). Red dots represent the best fitting model.

**Supplementary File 1:** zip-file containing the model and parameter estimation code (PEPSSBI), original Western blot images and quantified data, tables specifying all parameters, parameters bounds, and parameter estimates (Excel), model analysis code (Matlab), BMRI code (Matlab). Description of files:

Model code (folder): PEBSSPI files (.sbc). One file for each parameter estimation round.

Western blot images originals (folder): Bitmap images (.png) of the Western blots.

Data (Folder): Text files (.txt). Quantified, normalised Western blot data as text files.

BMRI (folder): Matlab code (.m, .mat). Code for Bayesian Modular Response analysis.

WB Quantification (Excel file): Tables with the quantified Western blot data.

Parameter bounds and estimation results (Excel file): Tables specifying the parameters and their estimation bounds and the results for each parameter estimation round. One sheet for each round.

Table model equations (Word file): Table specifying the model equations, including reactions, rate laws, algebraic constraints and ordinary differential equations.

