## Supplementary material for "Model-based identification of the crosstalks and feedbacks that determine the doxorubicin response dynamics of the JNK-p38-p53 network": Main and Supplementary Figures

Figure 1

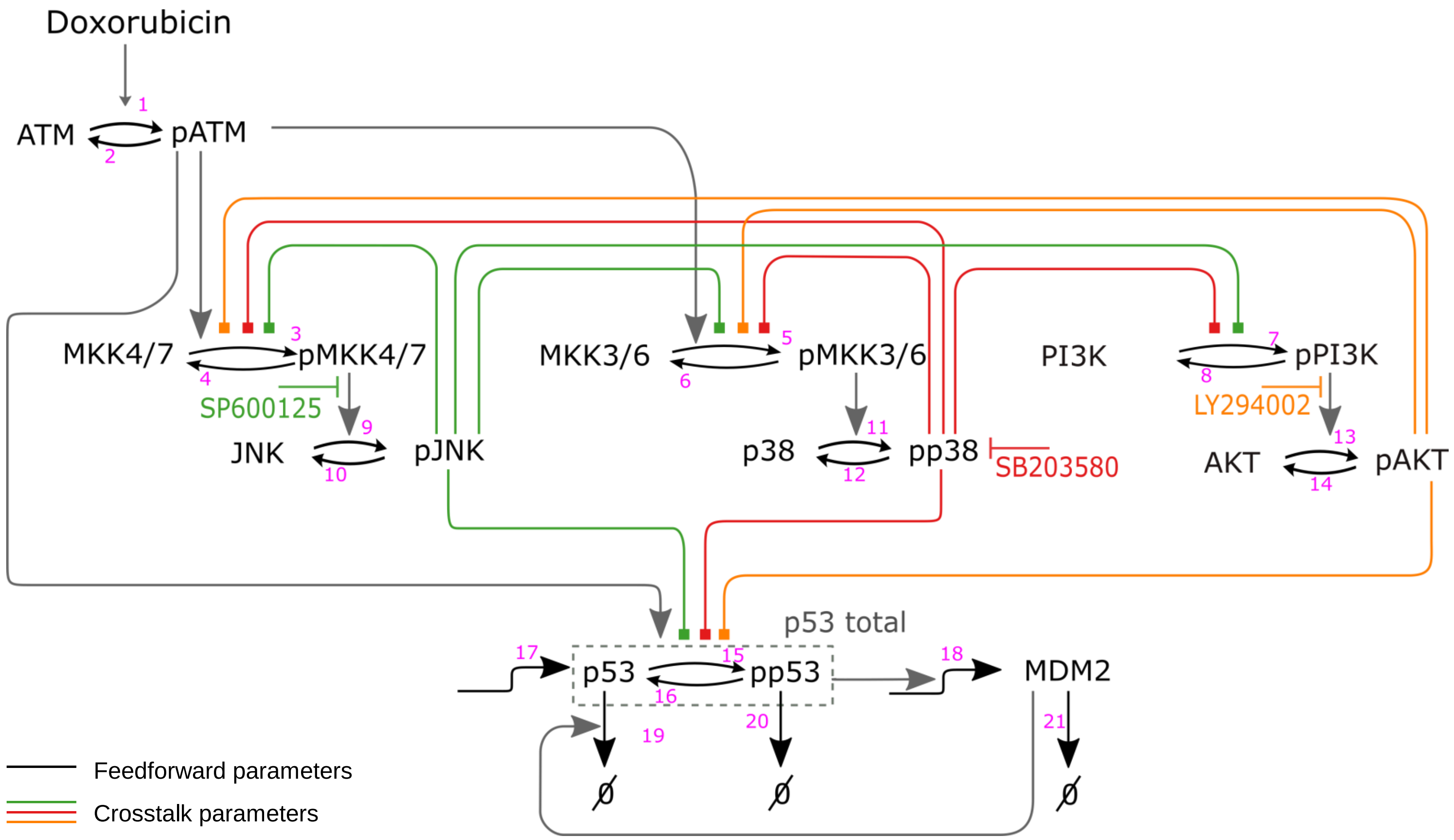

Doxorubicin
1
ATM
pATM
2
3
MKK4/7
pMKK4/7
4
SP600125
9
JNK
pJNK
10
5
MKK3/6
pMKK3/6
6
11
p38
pp38
12
SB203580
PI3K
7
pPI3K
8
LY294002
13
AKT
pAKT
14
p53 total
17
p53
15
pp53
16
18
MDM2
19
20
21
Ø
Ø
Ø
Feedforward parameters
Crosstalk parameters

Figure 2

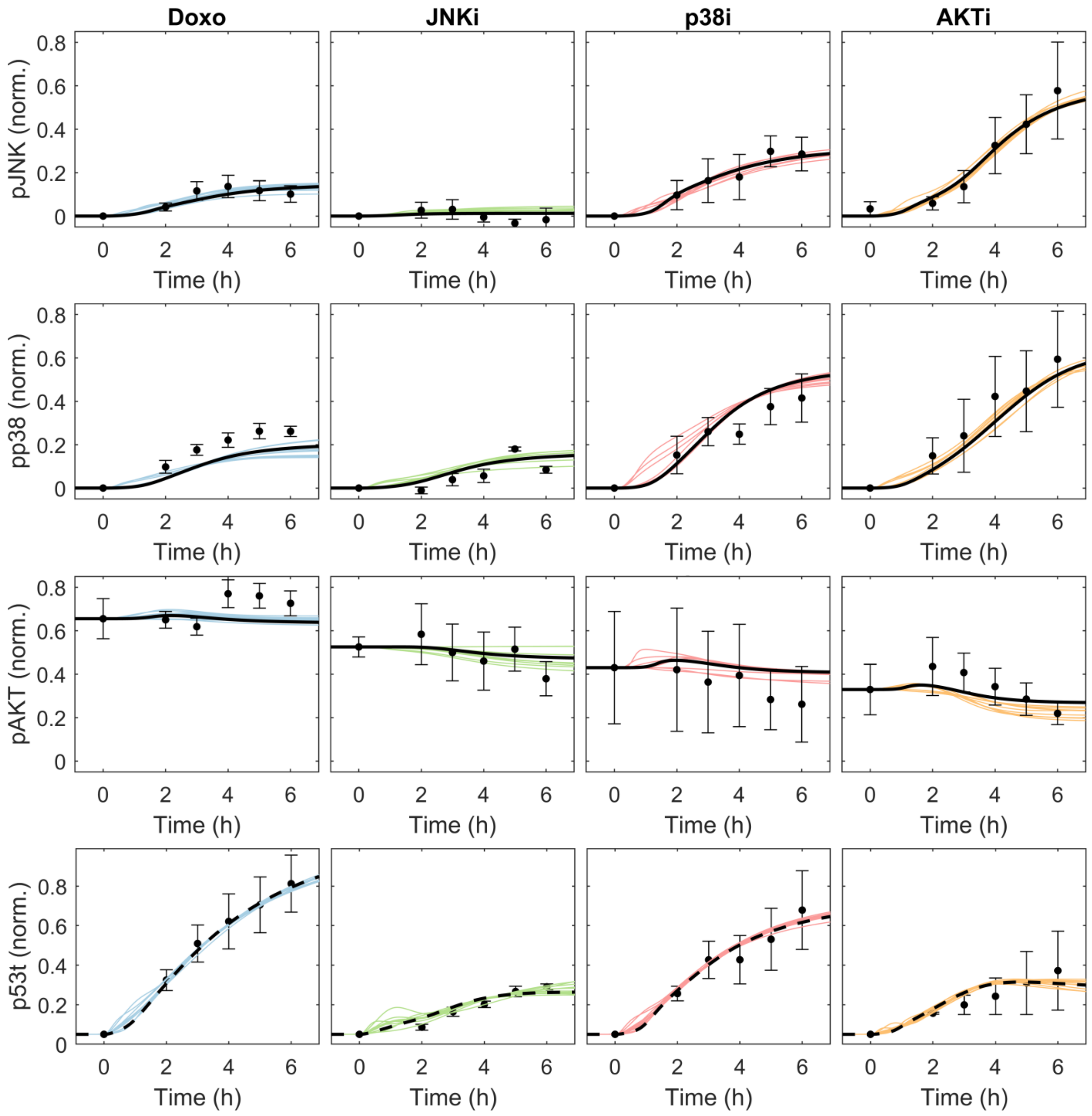
Doxo
JNKi
p38i
AKTi
pJNK (norm.)
pp38 (norm.)
pAKT (norm.)
p53t (norm.)
Time (h)

Figure 3

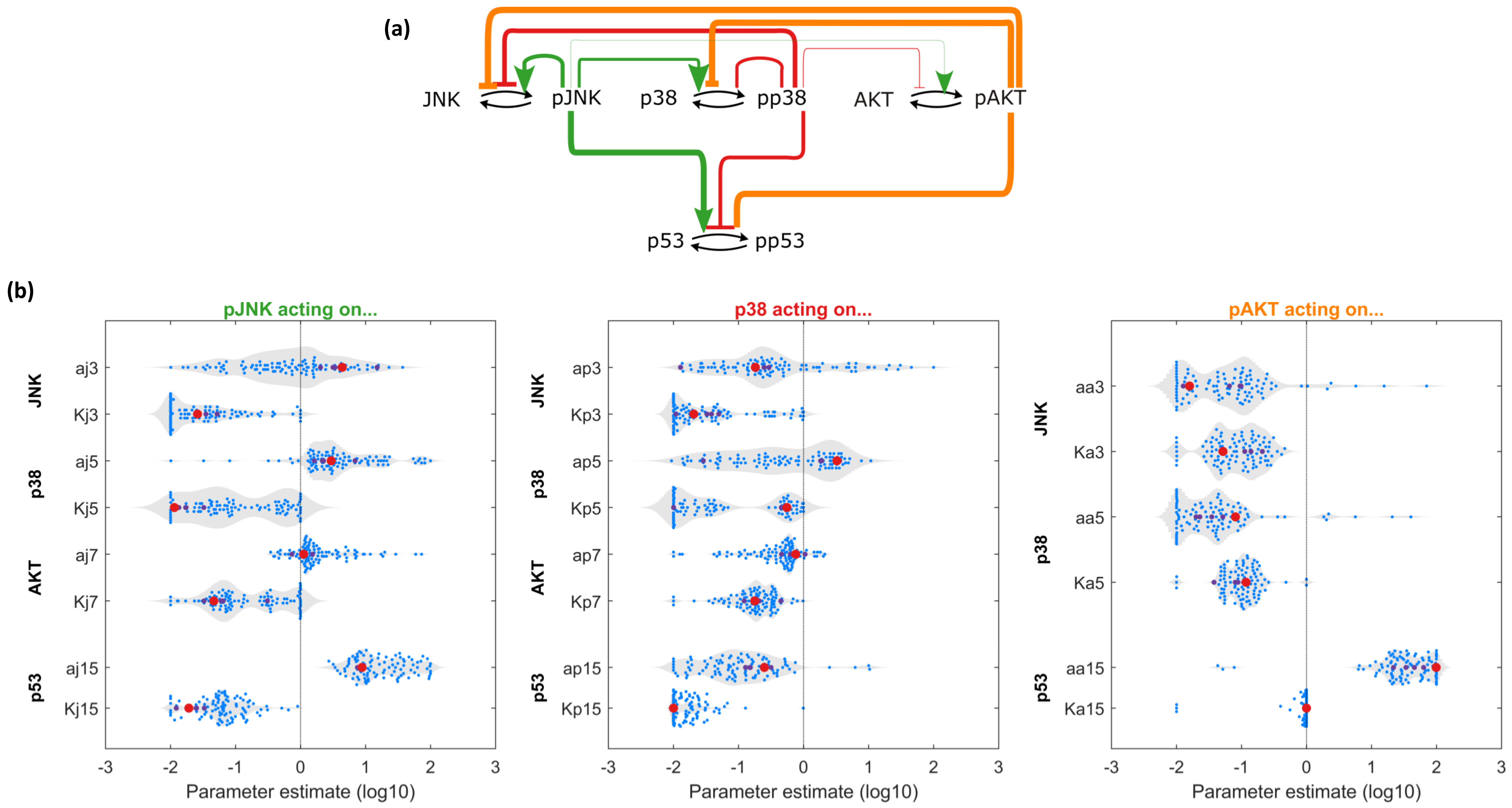
(a)
JNK
pJNK
p38
pp38
AKT
pAKT
p53
pp53
(b)
pJNK acting on...
p38 acting on...
pAKT acting on...
JNK
p38
AKT
p53
aj3
Kj3
aj5
Kj5
aj7
Kj7
aj15
Kj15
ap3
Kp3
ap5
Kp5
ap7
Kp7
ap15
Kp15
aa3
Ka3
aa5
Ka5
aa15
Ka15
-3 -2 -1 0 1 2 3
Parameter estimate (log10)

Figure 4

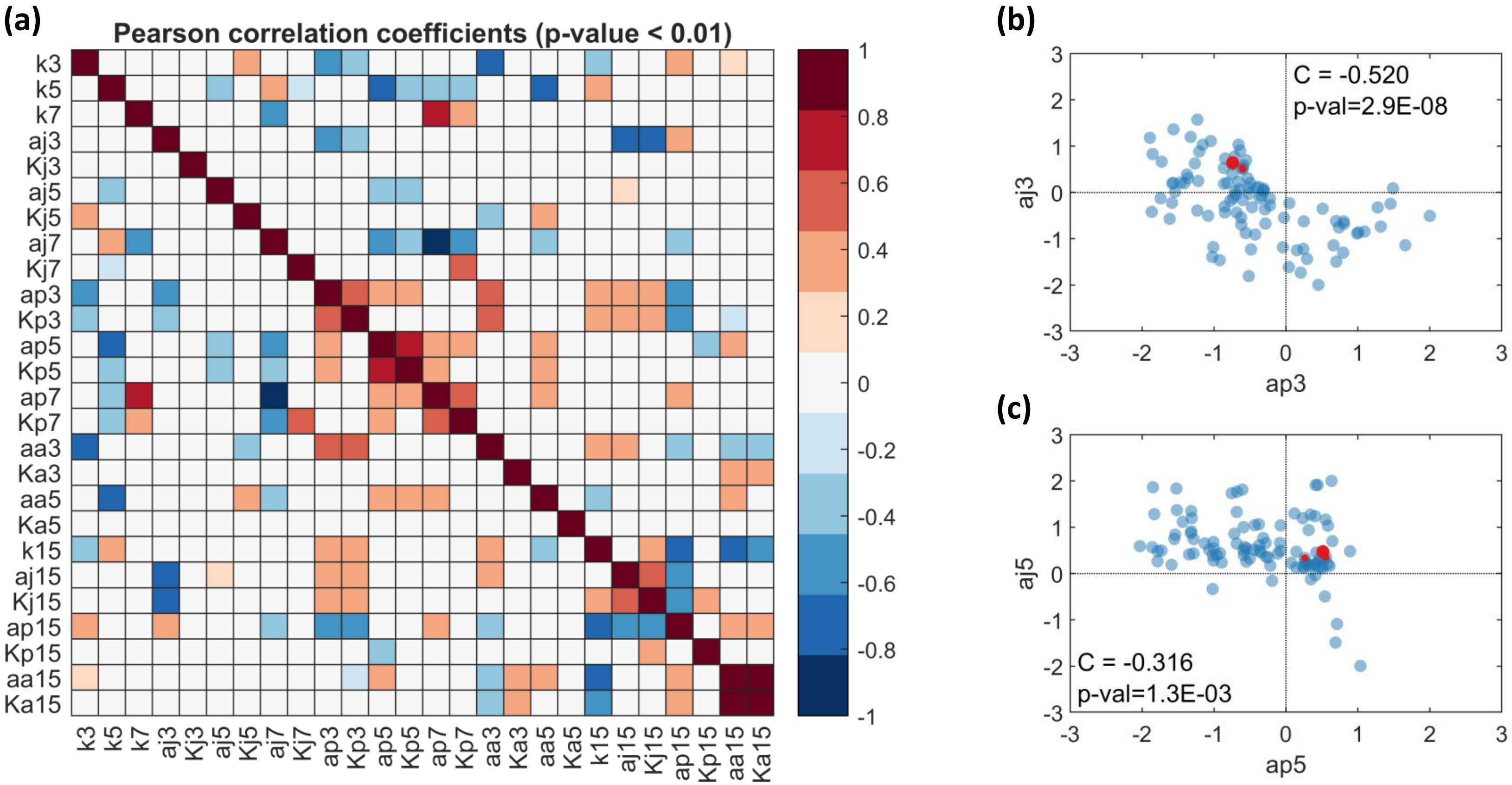
(a)
Pearson correlation coefficients (p-value < 0.01)
k3
k5
k7
aj3
Kj3
aj5
Kj5
aj7
Kj7
ap3
Kp3
ap5
Kp5
ap7
Kp7
aa3
Ka3
aa5
Ka5
k15
aj15
Kj15
ap15
Kp15
aa15
Ka15
1
0.8
0.6
0.4
0.2
0
-0.2
-0.4
-0.6
-0.8
-1
(b)
C = -0.520
p-val=2.9E-08
aj3
ap3
(c)
C = -0.316
p-val=1.3E-03
aj5
ap5

Figure 5

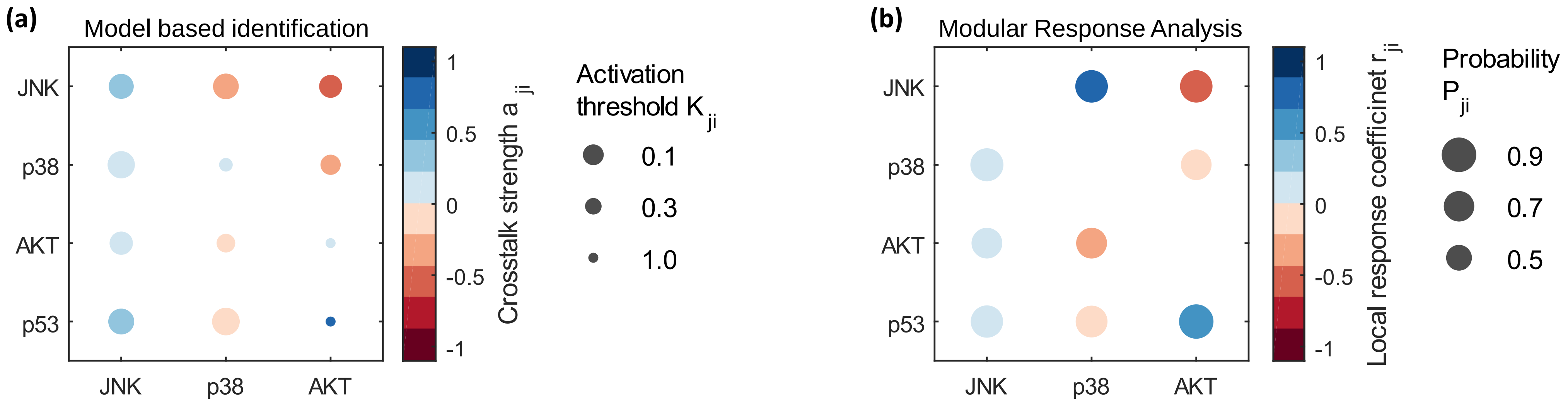

(a)
Model based identification
JNK
p38
AKT
p53
JNK
p38
AKT
1
0.5
0
-0.5
-1
Crosstalk strength a ji
Activation threshold K ji
0.1
0.3
1.0
(b)
Modular Response Analysis
JNK
p38
AKT
p53
JNK
p38
AKT
1
0.5
0
-0.5
-1
Local response coefficinet r ji
Probability P ji
0.9
0.7
0.5

Figure 6

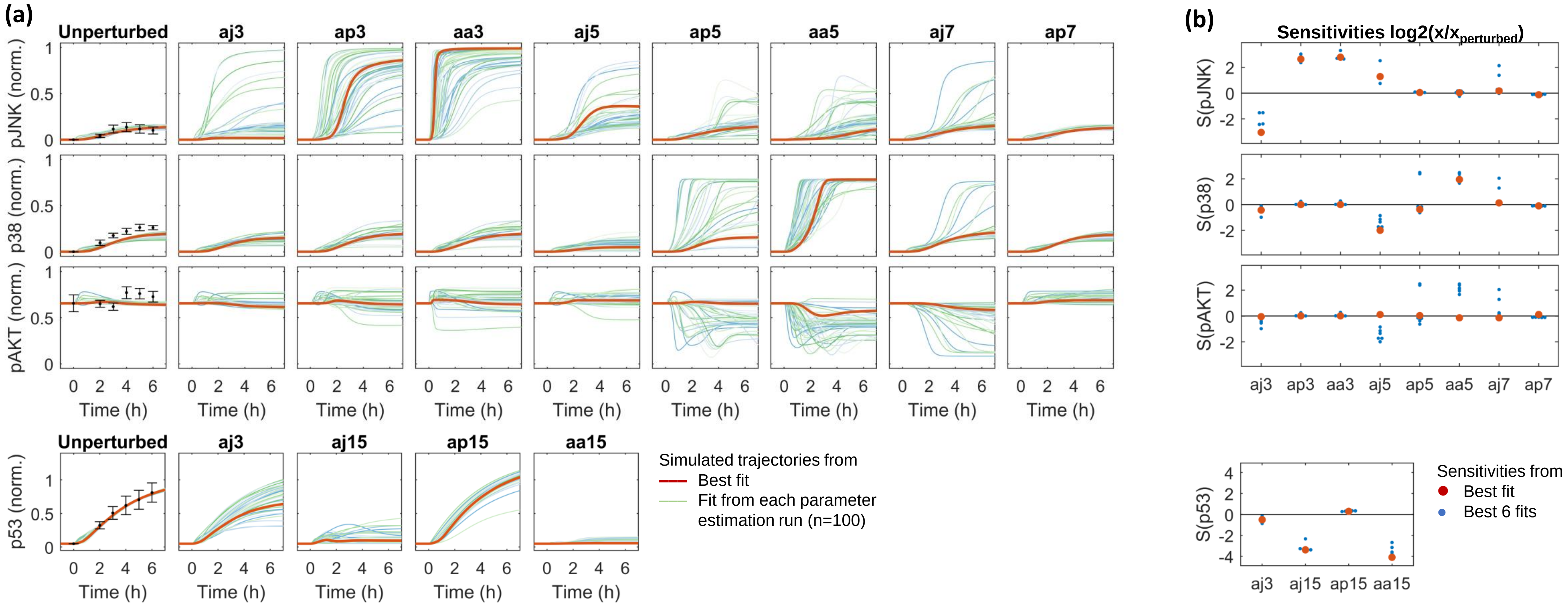

(a)
Unperturbed
aj3
ap3
aa3
aj5
ap5
aa5
aj7
ap7
pJNK (norm.)
p38 (norm.)
pAKT (norm.)
Time (h)
aj15
ap15
aa15
p53 (norm.)
Simulated trajectories from
Best fit
Fit from each parameter estimation run (n=100)
(b)
Sensitivities log2(x/x_perturbed)
S(pJNK)
S(p38)
S(pAKT)
S(p53)
Sensitivities from
Best fit
Best 6 fits

Figure S1

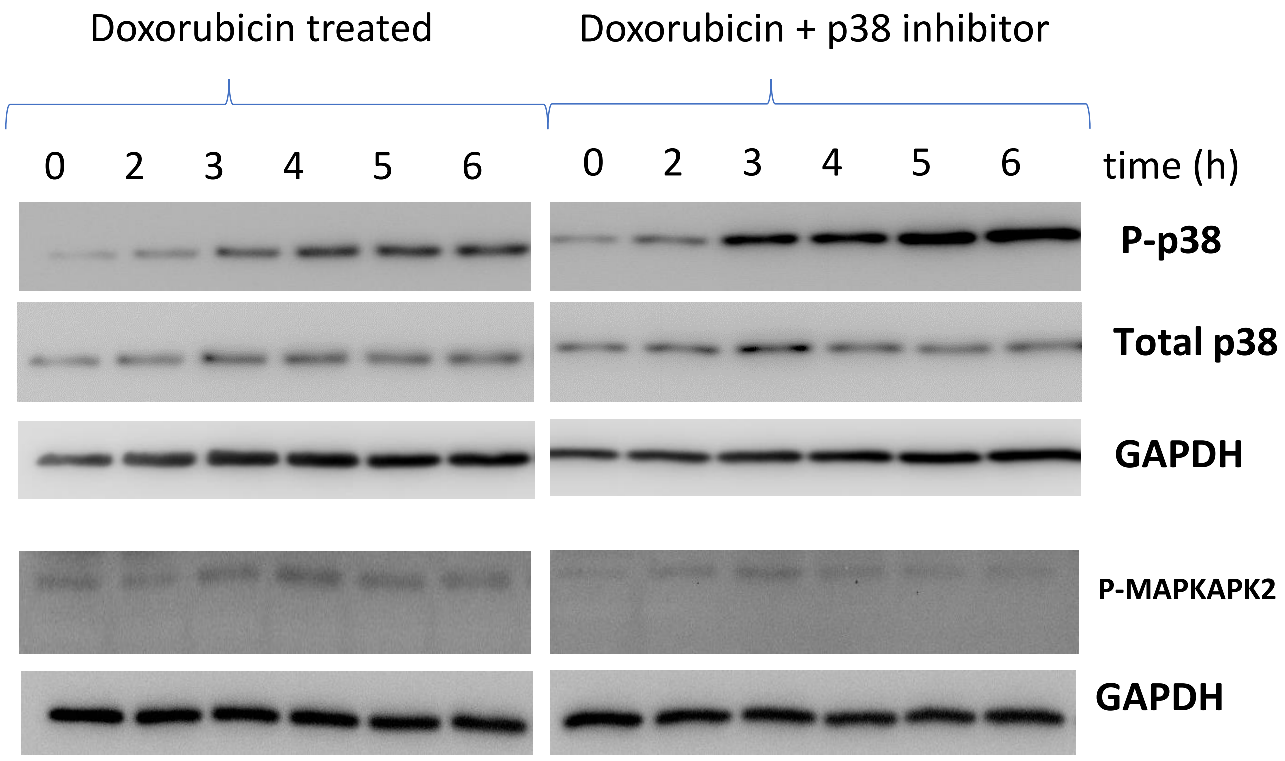


**Replicate 1**

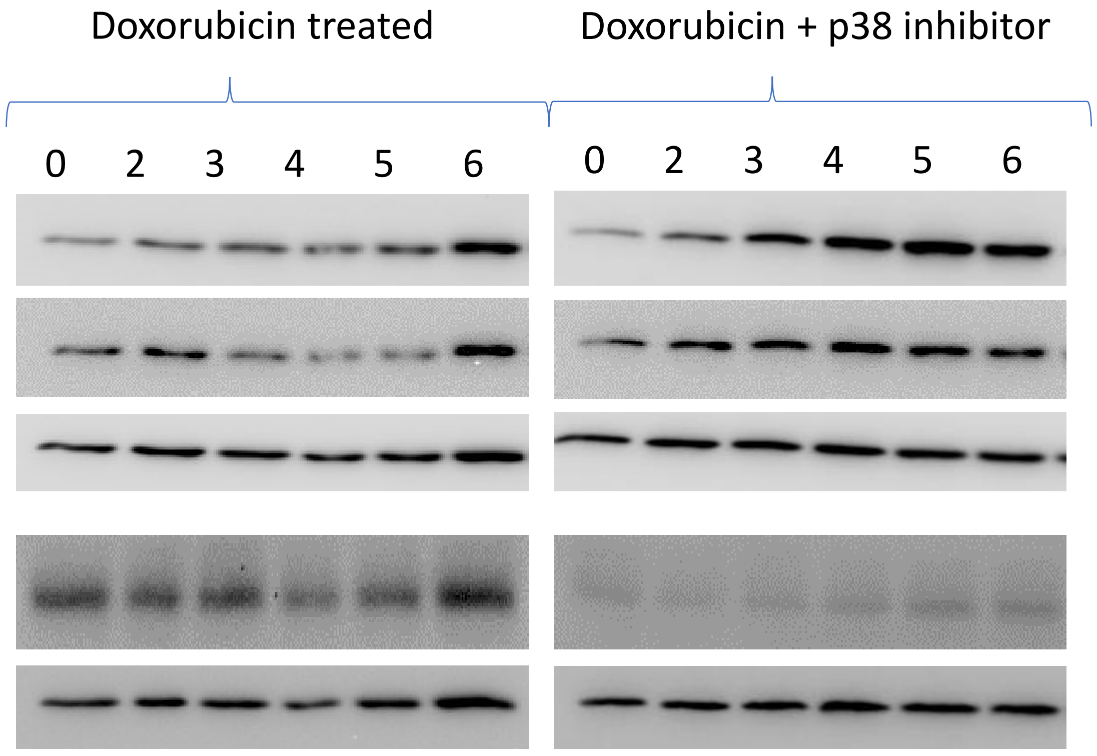

**Replicate 2**

Figure S2

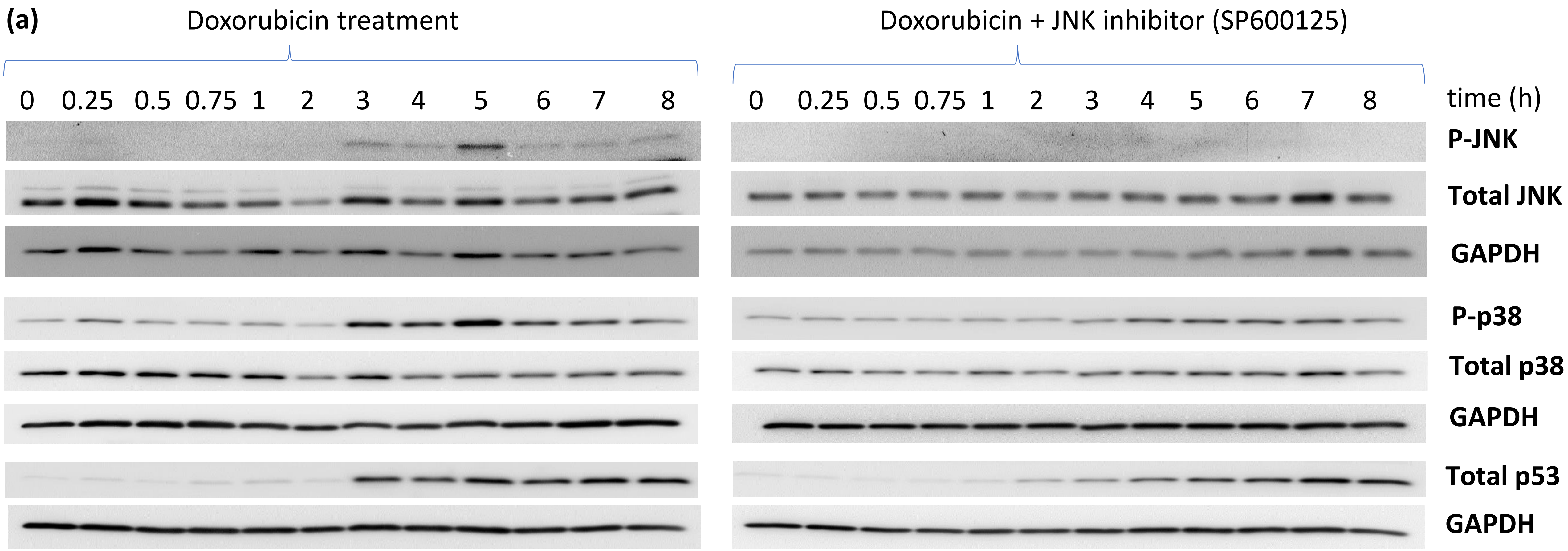
(a)
Doxorubicin treatment
Doxorubicin + JNK inhibitor (SP600125)
0 0.25 0.5 0.75 1 2 3 4 5 6 7 8
0 0.25 0.5 0.75 1 2 3 4 5 6 7 8
time (h)
P-JNK
Total JNK
GAPDH
P-p38
Total p38
GAPDH
Total p53
GAPDH

Figure S2

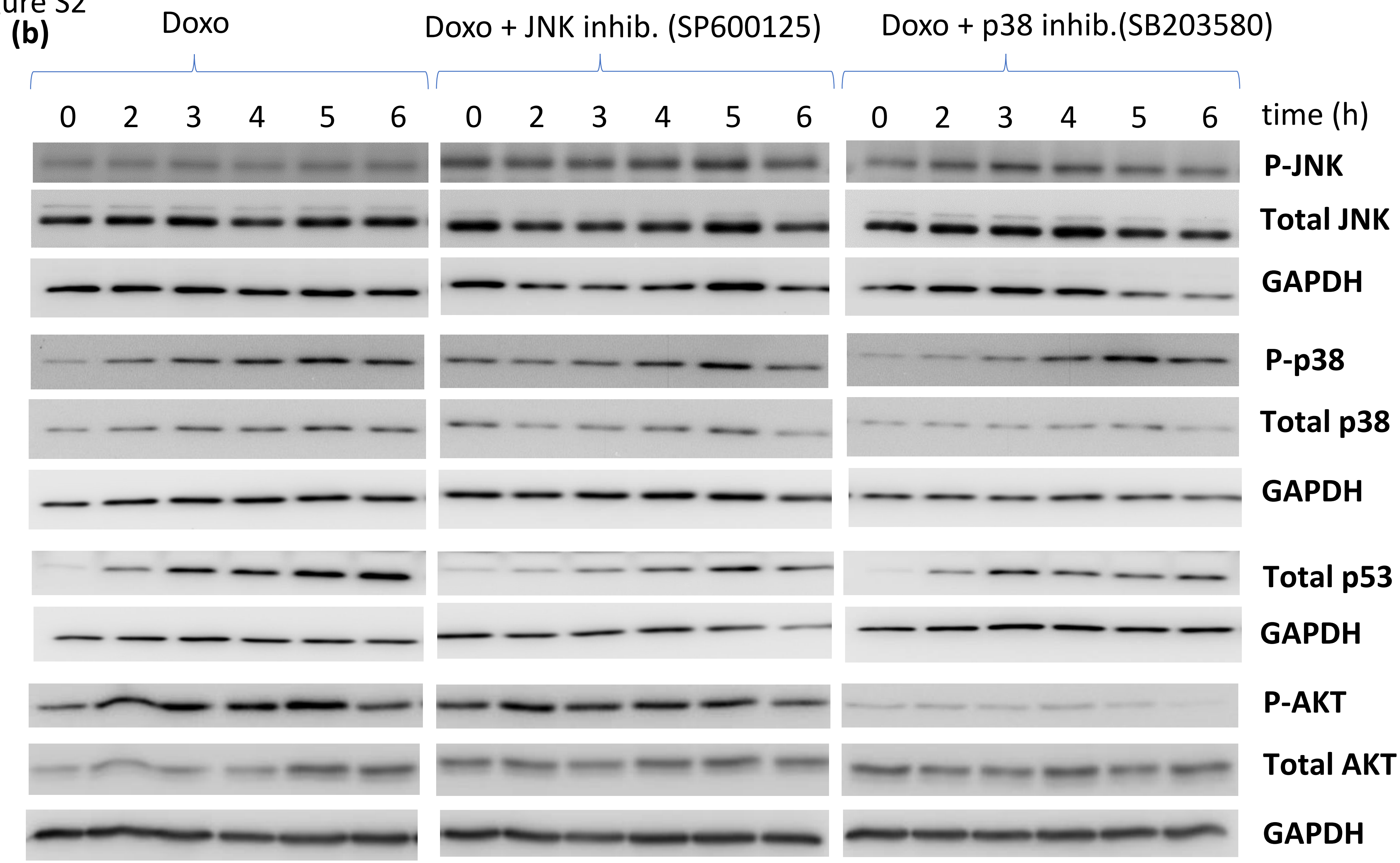
(b)
Doxo
Doxo + JNK inhib. (SP600125)
Doxo + p38 inhib.(SB203580)
0 2 3 4 5 6
0 2 3 4 5 6
0 2 3 4 5 6
time (h)
P-JNK
Total JNK
GAPDH
P-p38
Total p38
GAPDH
Total p53
GAPDH
P-AKT
Total AKT
GAPDH

Figure S2

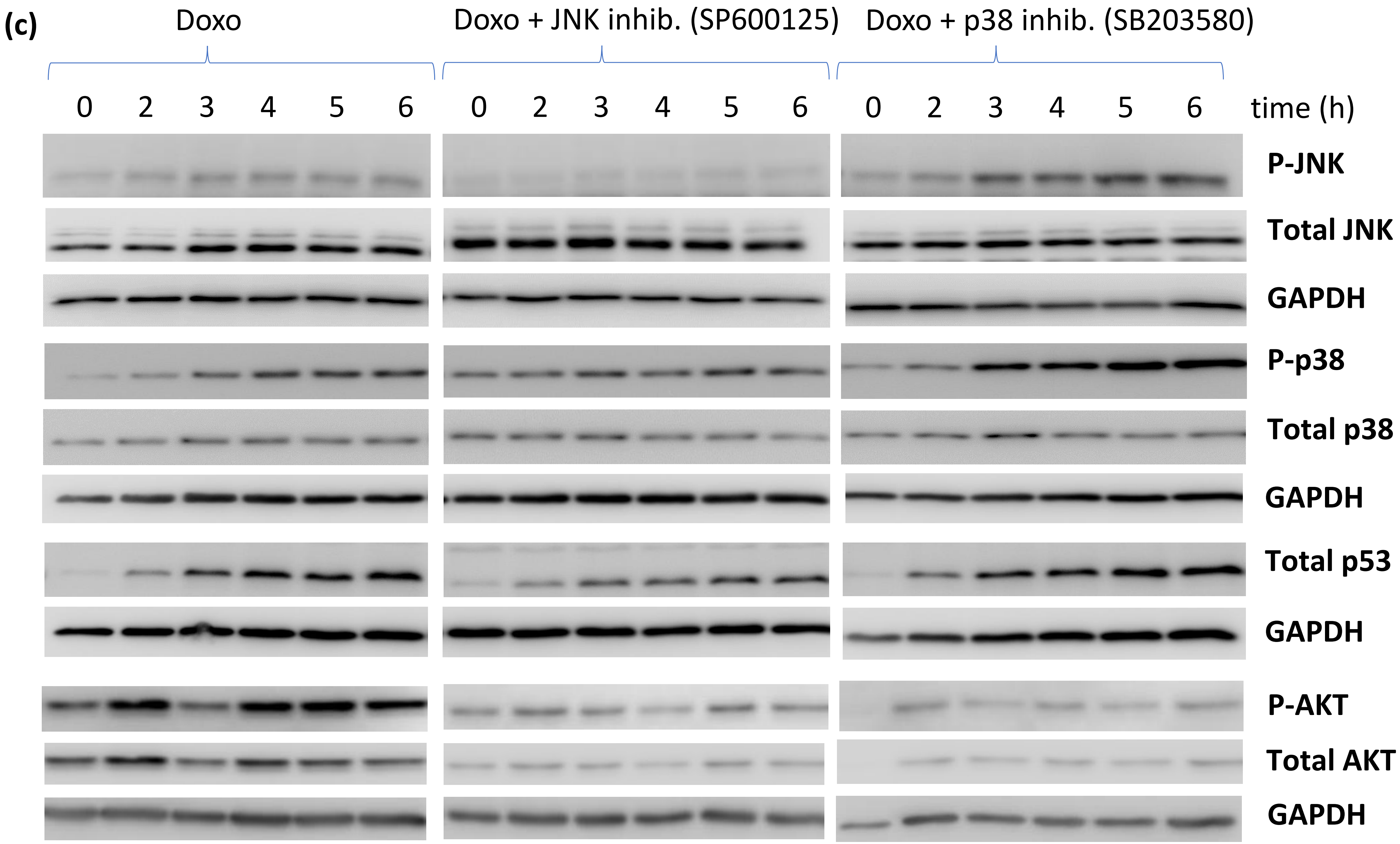
(c)
Doxo
Doxo + JNK inhib. (SP600125)
Doxo + p38 inhib. (SB203580)
0 2 3 4 5 6
0 2 3 4 5 6
0 2 3 4 5 6
time (h)
P-JNK
Total JNK
GAPDH
P-p38
Total p38
GAPDH
Total p53
GAPDH
P-AKT
Total AKT
GAPDH

Figure S2

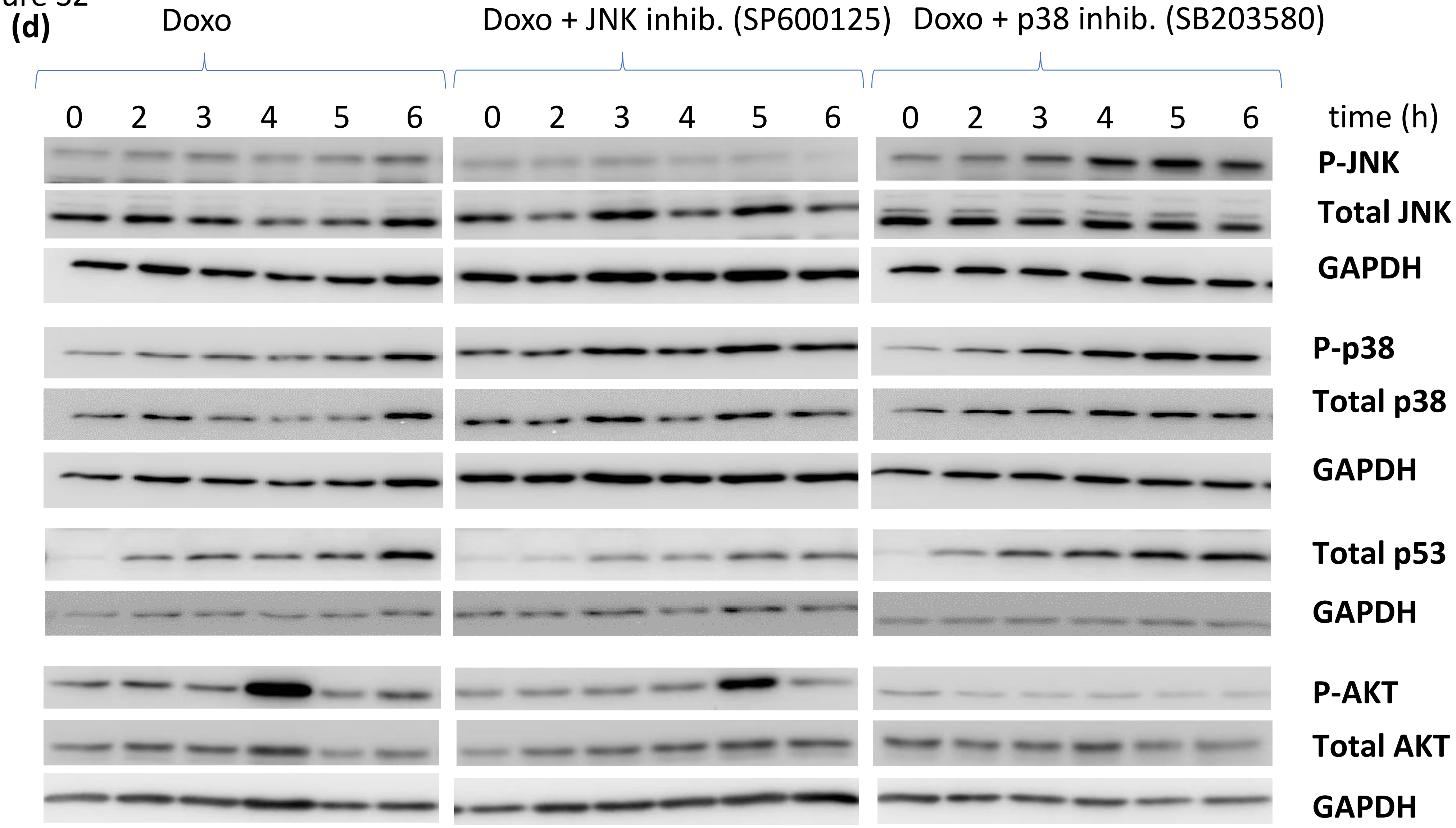
(d)
Doxo
Doxo + JNK inhib. (SP600125)
Doxo + p38 inhib. (SB203580)
0 2 3 4 5 6
0 2 3 4 5 6
0 2 3 4 5 6
time (h)
P-JNK
Total JNK
GAPDH
P-p38
Total p38
GAPDH
Total p53
GAPDH
P-AKT
Total AKT
GAPDH

Figure S2

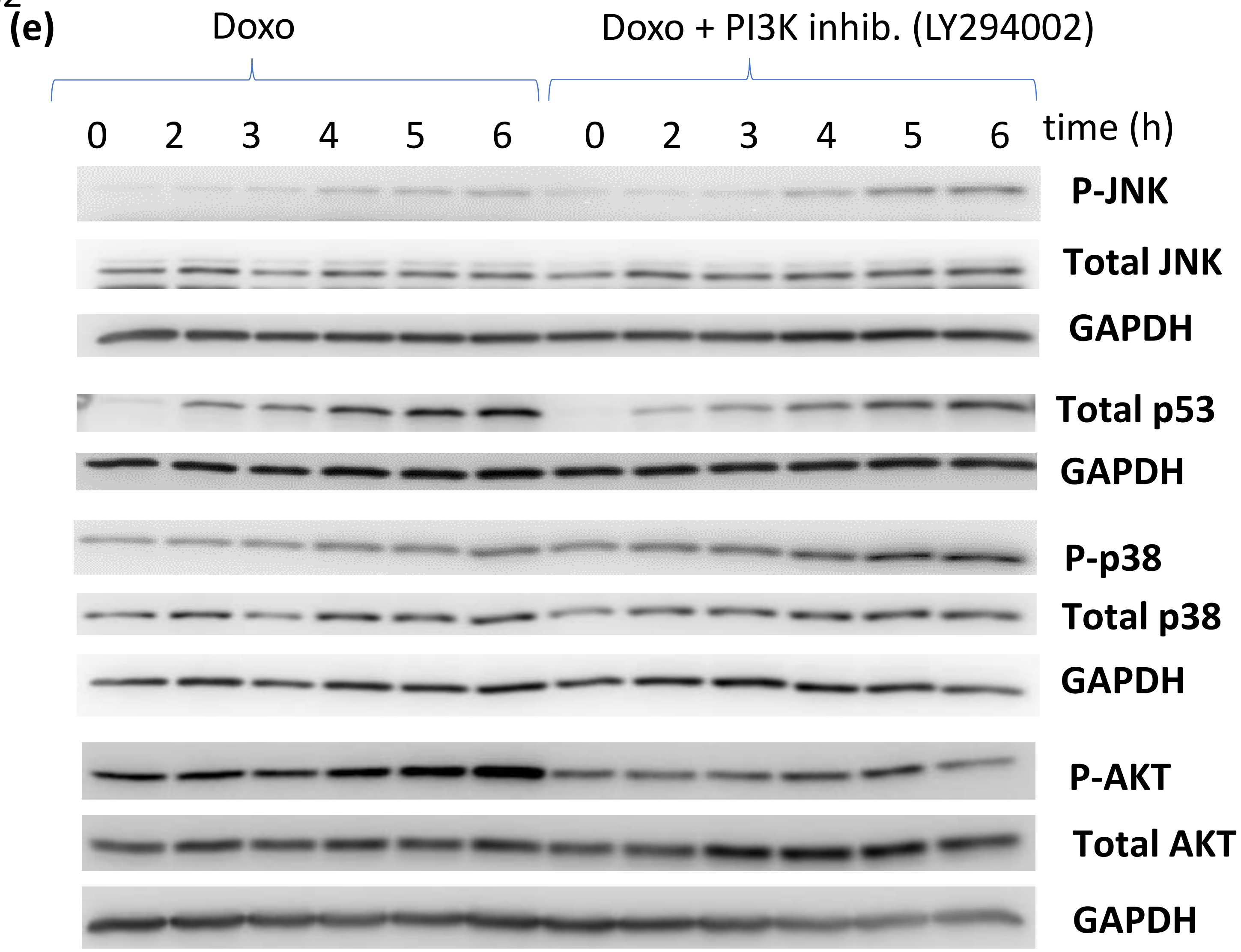
(e)
Doxo
Doxo + PI3K inhib. (LY294002)
0 2 3 4 5 6 0 2 3 4 5 6
time (h)
P-JNK
Total JNK
GAPDH
Total p53
GAPDH
P-p38
Total p38
GAPDH
P-AKT
Total AKT
GAPDH

Figure S2

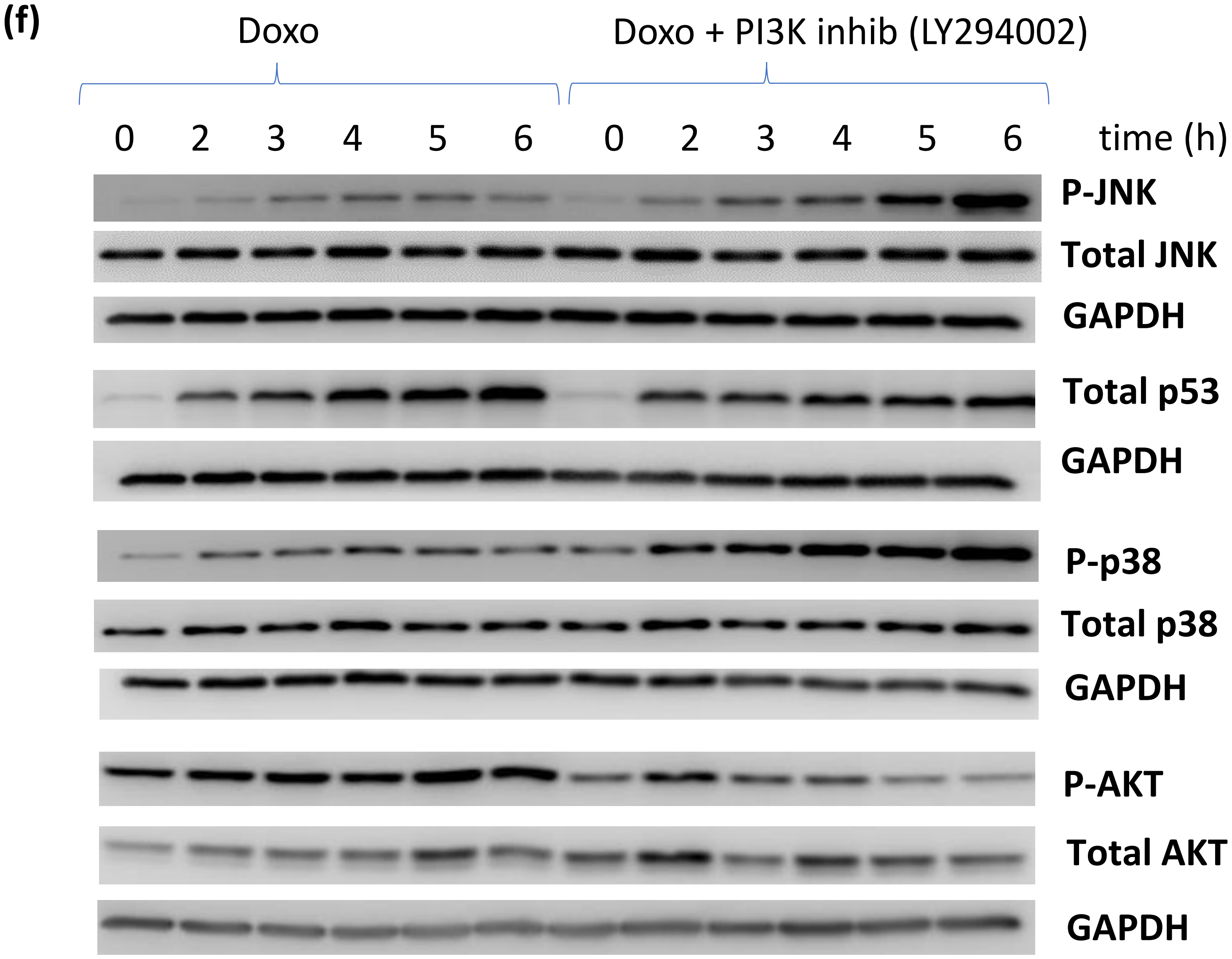
(f)
Doxo
Doxo + PI3K inhib (LY294002)
0 2 3 4 5 6 0 2 3 4 5 6 time (h)
P-JNK
Total JNK
GAPDH
Total p53
GAPDH
P-p38
Total p38
GAPDH
P-AKT
Total AKT
GAPDH

Figure S2

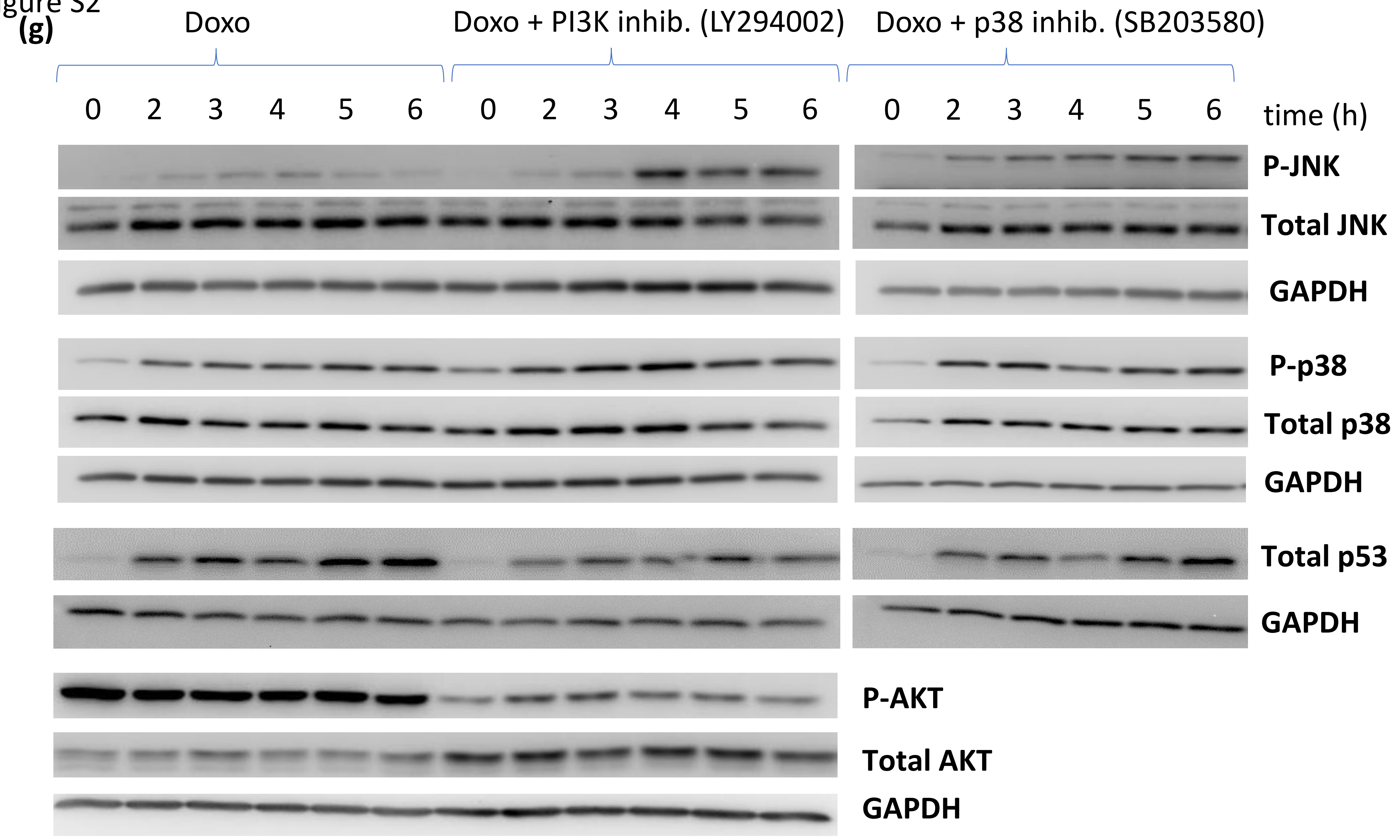
(g)
Doxo
Doxo + PI3K inhib. (LY294002)
Doxo + p38 inhib. (SB203580)
0 2 3 4 5 6 0 2 3 4 5 6 0 2 3 4 5 6 time (h)
P-JNK
Total JNK
GAPDH
P-p38
Total p38
GAPDH
Total p53
GAPDH
P-AKT
Total AKT
GAPDH

### Figure S3

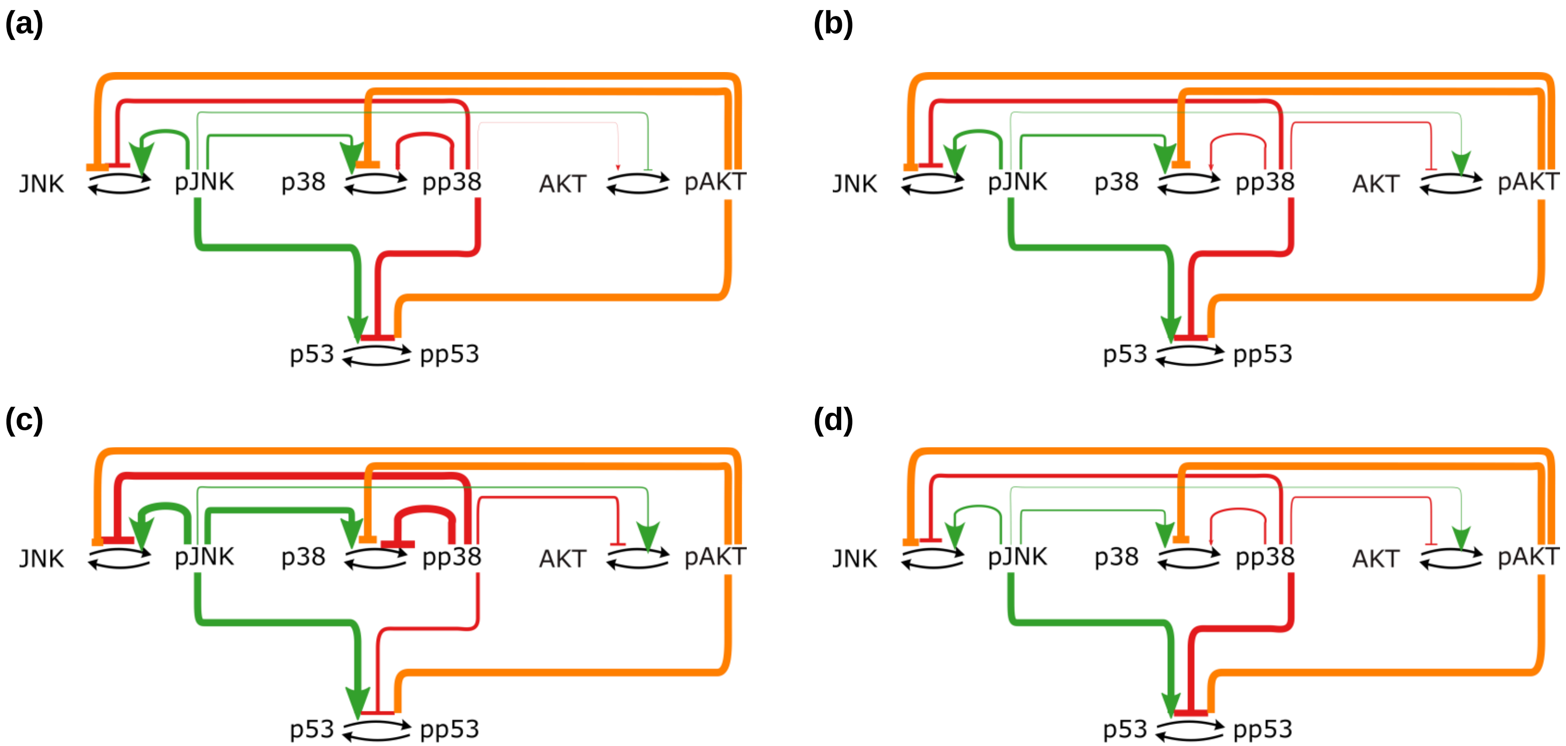
(a)
JNK
pJNK
p38
pp38
AKT
pAKT
p53
pp53
(b)
JNK
pJNK
p38
pp38
AKT
pAKT
p53
pp53
(c)
JNK
pJNK
p38
pp38
AKT
pAKT
p53
pp53
(d)
JNK
pJNK
p38
pp38
AKT
pAKT
p53
pp53

Figure S4

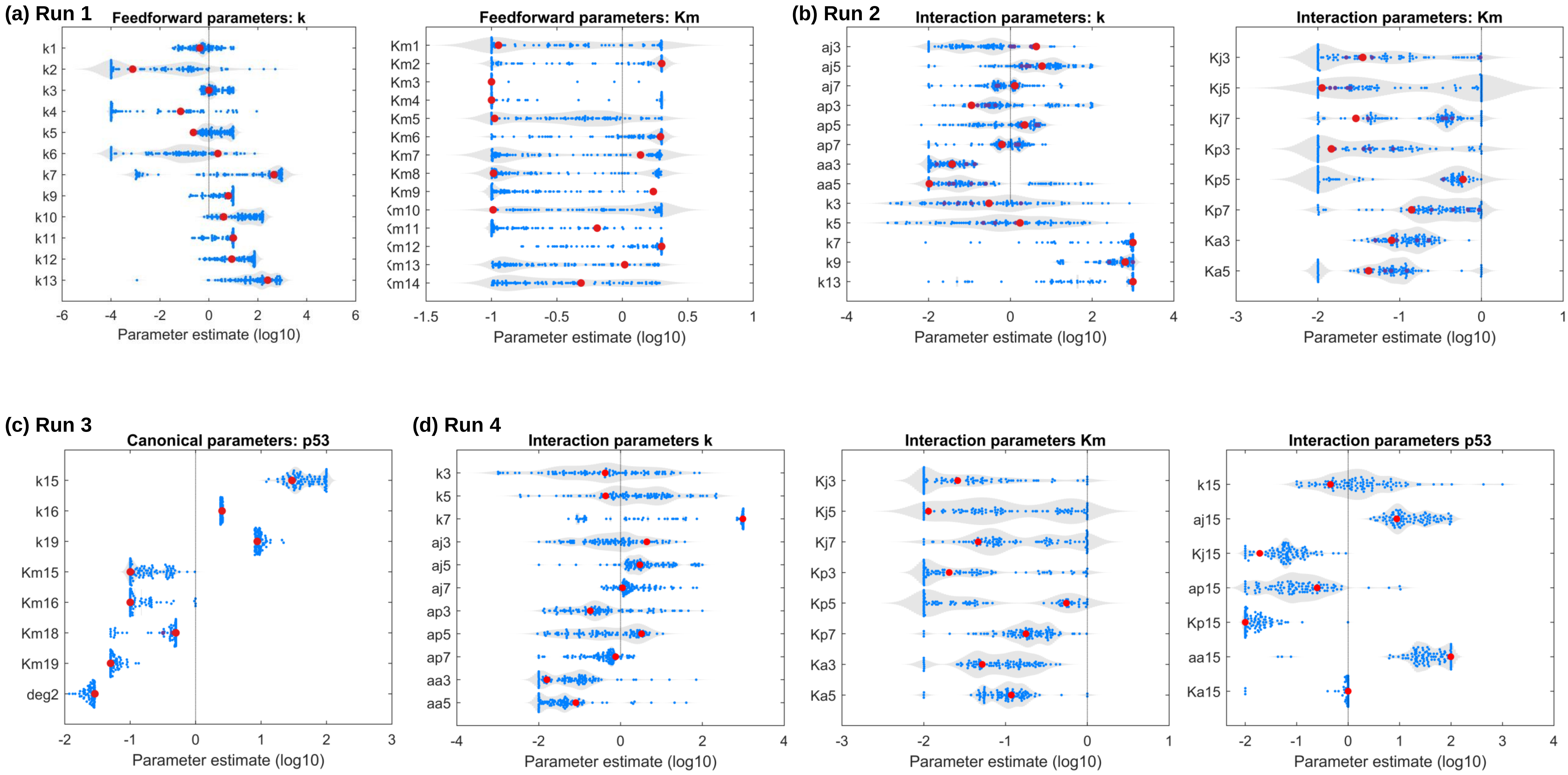

(a) Run 1
Feedforward parameters: k
k1
k2
k3
k4
k5
k6
k7
k9
k10
k11
k12
k13
-6
-4
-2
0
2
4
6
Parameter estimate (log10)
Feedforward parameters: Km
Km1
Km2
Km3
Km4
Km5
Km6
Km7
Km8
Km9
-1.5
-1
-0.5
0
0.5
1
Parameter estimate (log10)
(b) Run 2
Interaction parameters: k
aj3
aj5
aj7
ap3
ap5
ap7
aa3
aa5
k3
k5
k7
k9
k13
-4
-2
0
2
4
Parameter estimate (log10)
Interaction parameters: Km
Kj3
Kj5
Kj7
Kp3
Kp5
Kp7
Ka3
Ka5
-3
-2
-1
0
1
Parameter estimate (log10)
(c) Run 3
Canonical parameters: p53
k15
k16
k19
Km15
Km16
Km18
Km19
deg2
-2
-1
0
1
2
3
Parameter estimate (log10)
(d) Run 4
Interaction parameters k
k3
k5
k7
aj3
aj5
aj7
ap3
ap5
ap7
aa3
aa5
-4
-2
0
2
4
Parameter estimate (log10)
Interaction parameters Km
Kj3
Kj5
Kj7
Kp3
Kp5
Kp7
Ka3
Ka5
-3
-2
-1
0
1
Parameter estimate (log10)
Interaction parameters p53
k15
aj15
Kj15
ap15
Kp15
aa15
Ka15
-2
-1
0
1
2
3
4
Parameter estimate (log10)

### Figure S5

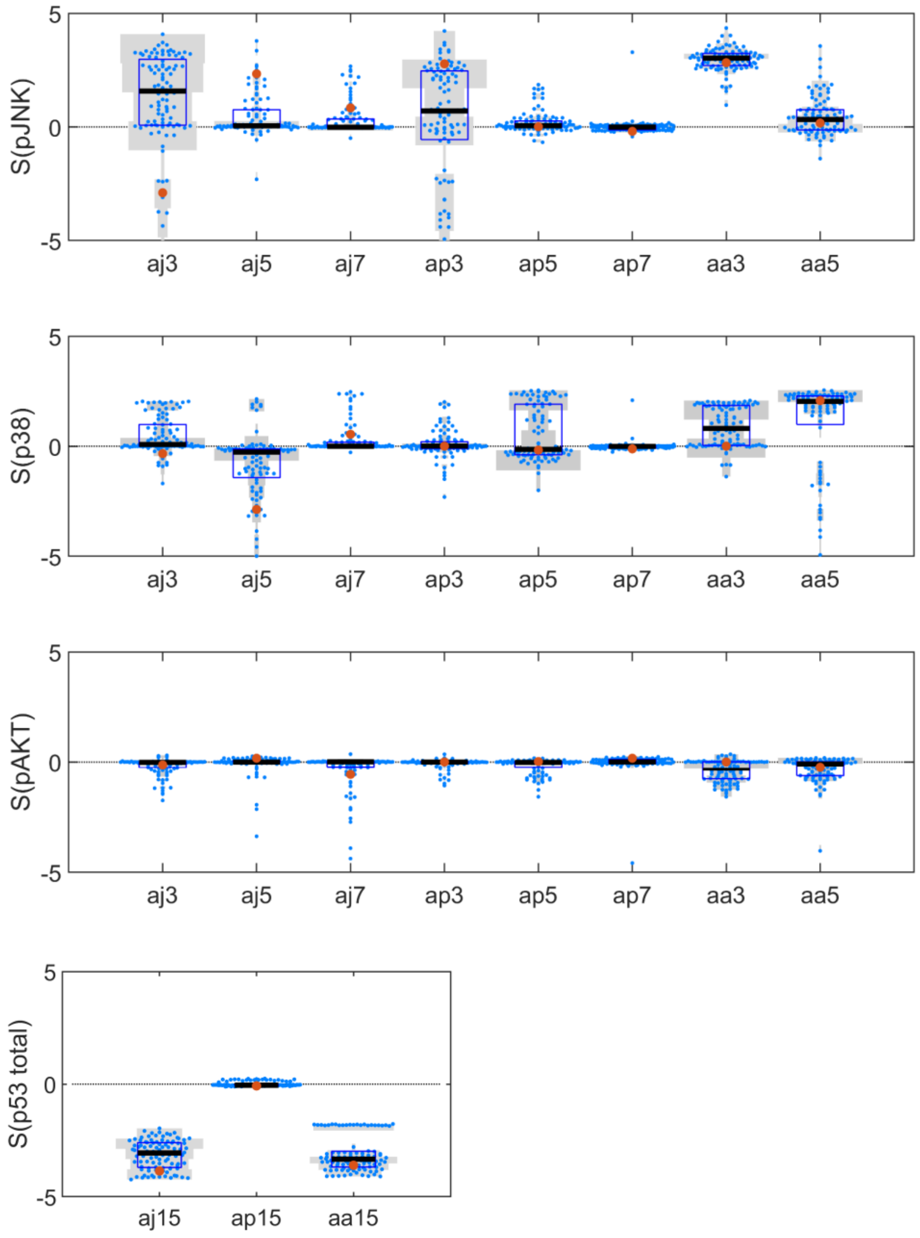

S(pJNK)
S(p38)
S(pAKT)
S(p53 total)
5
0
-5
aj3
aj5
aj7
ap3
ap5
ap7
aa3
aa5
aj15
ap15
aa15
