## Supplementary material for "Model-based identification of the crosstalks and feedbacks that determine the doxorubicin response dynamics of the JNK-p38-p53 network": Table model equations

| Table 1: List of biochemical reactions, associated rate laws, algebraic constraints and differential equations | | |
| --- | --- | --- |
| Reactions | Reaction equation | Cross talks and feedbacks |
| i) Upstream pathways |  |  |
| ATM $\underset{\to}{\boldsymbol{d}\boldsymbol{oxo}}$ pATM | $\boldsymbol{v}\boldsymbol{1=k}\boldsymbol{1*doxo*}\frac{\boldsymbol{ATM}}{\boldsymbol{ATM+Km}\boldsymbol{1}}\boldsymbol{,}$ | where $\boldsymbol{doxo}=\left\{ \begin{aligned} 0 \text{for} t<0 \\ 1 \text{for} t\geq0 \end{aligned} \right.$ |
| associated reverse reaction | $\boldsymbol{v2=k}\boldsymbol{2*}\frac{\boldsymbol{pATM}}{\boldsymbol{pATM+Km}\boldsymbol{2}}$ |  |
| MKK4/7 $\underset{\to}{\boldsymbol{p}\boldsymbol{ATM}}$ pMKK4/7 | $\boldsymbol{v}\boldsymbol{3=k}\boldsymbol{3*pATM*}\frac{\boldsymbol{MKK}\boldsymbol{4/7}}{\boldsymbol{MKK}\boldsymbol{4/7+Km}\boldsymbol{3}}$ | $\boldsymbol{*}{\frac{{\boldsymbol{Kp}\boldsymbol{3}}^{\boldsymbol{np}\boldsymbol{3}}\boldsymbol{+ap}\boldsymbol{3*ap}\boldsymbol{38}}{{\boldsymbol{Kp}\boldsymbol{3}}^{\boldsymbol{np}\boldsymbol{3}}\boldsymbol{+}{\boldsymbol{ap}\boldsymbol{38}}^{\boldsymbol{np}\boldsymbol{3}}}}^{\boldsymbol{np}\boldsymbol{3}}\boldsymbol{*}{\frac{{\boldsymbol{Ka}\boldsymbol{3}}^{\boldsymbol{na}\boldsymbol{3}}\boldsymbol{+aa3*pAKT}}{{\boldsymbol{Ka}\boldsymbol{3}}^{\boldsymbol{na}\boldsymbol{3}}\boldsymbol{+}\boldsymbol{pAKT}^{\boldsymbol{na}\boldsymbol{3}}}}^{\boldsymbol{na}\boldsymbol{3}}\boldsymbol{*}{\frac{{\boldsymbol{Kj}\boldsymbol{3}}^{\boldsymbol{nj}\boldsymbol{3}}\boldsymbol{+aj3*pJNK}}{{\boldsymbol{Kj}\boldsymbol{3}}^{\boldsymbol{nj}\boldsymbol{3}}\boldsymbol{+}\boldsymbol{pJNK}^{\boldsymbol{nj}\boldsymbol{3}}}}^{\boldsymbol{nj}\boldsymbol{3}}$ |
| associated reverse reaction | $\boldsymbol{v}\boldsymbol{4=k}\boldsymbol{4*}\frac{\boldsymbol{pMKK}\boldsymbol{4/7}}{\boldsymbol{pMKK}\boldsymbol{4/7+Km}\boldsymbol{4}}$ |  |
| MKK3/6 $\underset{\to}{\boldsymbol{p}\boldsymbol{ATM}}$ pMKK3/6 | $\boldsymbol{v}\boldsymbol{5=k}\boldsymbol{5*pATM*}\frac{\boldsymbol{MKK}\boldsymbol{3/6}}{\boldsymbol{MKK}\boldsymbol{3/6+Km}\boldsymbol{5}}$ | $\boldsymbol{*}{\frac{{\boldsymbol{Kj}\boldsymbol{5}}^{\boldsymbol{nj}\boldsymbol{5}}\boldsymbol{+aj}\boldsymbol{5*pJNK}}{{\boldsymbol{Kj}\boldsymbol{5}}^{\boldsymbol{nj}\boldsymbol{5}}\boldsymbol{+}\boldsymbol{pJNK}^{\boldsymbol{nj}\boldsymbol{5}}}}^{\boldsymbol{nj}\boldsymbol{5}}\boldsymbol{*}{\frac{{\boldsymbol{Ka}\boldsymbol{5}}^{\boldsymbol{na}\boldsymbol{5}}\boldsymbol{+aa}\boldsymbol{5*pAKT}}{{\boldsymbol{Ka}\boldsymbol{5}}^{\boldsymbol{na}\boldsymbol{5}}\boldsymbol{+}\boldsymbol{pAKT}^{\boldsymbol{na}\boldsymbol{5}}}}^{\boldsymbol{na}\boldsymbol{5}}\boldsymbol{*}{\frac{{\boldsymbol{Kp}\boldsymbol{5}}^{\boldsymbol{np}\boldsymbol{5}}\boldsymbol{+ap}\boldsymbol{5*ap}\boldsymbol{38}}{{\boldsymbol{Kp}\boldsymbol{5}}^{\boldsymbol{np}\boldsymbol{5}}\boldsymbol{+}\boldsymbol{ap38}^{\boldsymbol{np}\boldsymbol{5}}}}^{\boldsymbol{np}\boldsymbol{5}}$ |
| associated reverse reaction | $\boldsymbol{v}\boldsymbol{6=k}\boldsymbol{6*}\frac{\boldsymbol{pMKK}\boldsymbol{3/6}}{\boldsymbol{pMKK}\boldsymbol{3/6+Km}\boldsymbol{6}}$ |  |
| PI3K $\underset{\to}{}$ pPI3K | $\boldsymbol{v}\boldsymbol{7=k}\boldsymbol{7*}\frac{\boldsymbol{PI}\boldsymbol{3}\boldsymbol{K}}{\boldsymbol{PI}\boldsymbol{3}\boldsymbol{K+Km}\boldsymbol{7}}$ | $\boldsymbol{*}{\frac{{\boldsymbol{Kp}\boldsymbol{7}}^{\boldsymbol{np}\boldsymbol{7}}\boldsymbol{+ap}\boldsymbol{7*ap}\boldsymbol{38}}{{\boldsymbol{Kp}\boldsymbol{7}}^{\boldsymbol{np}\boldsymbol{7}}\boldsymbol{+}{\boldsymbol{ap}\boldsymbol{38}}^{\boldsymbol{np}\boldsymbol{7}}}}^{\boldsymbol{np}\boldsymbol{7}}\boldsymbol{*}{\frac{{\boldsymbol{Kj}\boldsymbol{7}}^{\boldsymbol{nj7}}\boldsymbol{+aj}\boldsymbol{7*pJNK}}{{\boldsymbol{Kj}\boldsymbol{7}}^{\boldsymbol{nj}\boldsymbol{7}}\boldsymbol{+}\boldsymbol{pJNK}^{\boldsymbol{nj}\boldsymbol{7}}}}^{\boldsymbol{nj}\boldsymbol{7}}$ |
| associated reverse reaction | $\boldsymbol{v}\boldsymbol{8=k}\boldsymbol{8*}\frac{\boldsymbol{pPI}\boldsymbol{3}\boldsymbol{K}}{\boldsymbol{pPI}\boldsymbol{3}\boldsymbol{K+Km}\boldsymbol{8}}$ |  |
| JNK $\underset{\to}{\boldsymbol{p}\boldsymbol{MKK}\boldsymbol{4/7}}$ pJNK | $\boldsymbol{v9=k}\boldsymbol{9*pMKK}\boldsymbol{4/7 *}\frac{\boldsymbol{JNK}}{\boldsymbol{JNK+Km}\boldsymbol{9}}\boldsymbol{* SP}$ | |
| associated reverse reaction | $\boldsymbol{v}\boldsymbol{10=k10*}\frac{\boldsymbol{pJNK}}{\boldsymbol{pJNK+Km}\boldsymbol{10}}$ |  |
| p38 $\underset{\to}{\boldsymbol{pMKK}\boldsymbol{3/6}}$ pp38 | $\boldsymbol{v}\boldsymbol{11=k}\boldsymbol{11*pMKK}\boldsymbol{3/6*}\frac{\boldsymbol{p}\boldsymbol{38}}{\boldsymbol{p}\boldsymbol{38+Km}\boldsymbol{11}}$ | |
| associated reverse reaction | $\boldsymbol{v}\boldsymbol{12=k}\boldsymbol{12*}\frac{\boldsymbol{pp}\boldsymbol{38}}{\boldsymbol{pp}\boldsymbol{38+Km}\boldsymbol{12}}$ |  |
| AKT $\underset{\to}{\boldsymbol{pPI}\boldsymbol{3}\boldsymbol{K}}$ pAKT | $\boldsymbol{v}\boldsymbol{13=k}\boldsymbol{13*pPI}\boldsymbol{3}\boldsymbol{K}\frac{\boldsymbol{AKT}}{\boldsymbol{AKT+Km}\boldsymbol{13}}$ $\boldsymbol{* LY}$ | |
| associated reverse reaction | $\boldsymbol{v}\boldsymbol{14=k}\boldsymbol{14*}\frac{\boldsymbol{pAKT}}{\boldsymbol{pAKT+Km}\boldsymbol{14}}$ |  |
| II) Downstream pathway |  |  |
| p53 $\underset{\to}{\boldsymbol{p}\boldsymbol{ATM}}$ pp53 | $\boldsymbol{v}\boldsymbol{15=k}\boldsymbol{15*pATM*}\frac{\boldsymbol{p}\boldsymbol{53}}{\boldsymbol{p53+Km}\boldsymbol{15}}\boldsymbol{*}{\frac{{\boldsymbol{Kp}\boldsymbol{15}}^{\boldsymbol{np}\boldsymbol{15}}\boldsymbol{+ap}\boldsymbol{15*ap}\boldsymbol{38}}{{\boldsymbol{Kp}\boldsymbol{15}}^{\boldsymbol{np}\boldsymbol{15}}\boldsymbol{+}{\boldsymbol{ap}\boldsymbol{38}}^{\boldsymbol{np}\boldsymbol{15}}}}^{\boldsymbol{np}\boldsymbol{15}}\boldsymbol{*}{\frac{{\boldsymbol{Kj}\boldsymbol{15}}^{\boldsymbol{nj}\boldsymbol{15}}\boldsymbol{+aj}\boldsymbol{15*pJNK}}{{\boldsymbol{Kj}\boldsymbol{15}}^{\boldsymbol{nj}\boldsymbol{15}}\boldsymbol{+}\boldsymbol{pJNK}^{\boldsymbol{nj}\boldsymbol{15}}}}^{\boldsymbol{nj}\boldsymbol{15}}\boldsymbol{*}{\frac{{\boldsymbol{Ka}\boldsymbol{15}}^{\boldsymbol{na}\boldsymbol{15}}\boldsymbol{+aa}\boldsymbol{15*pAKT}}{{\boldsymbol{Ka}\boldsymbol{15}}^{\boldsymbol{na}\boldsymbol{15}}\boldsymbol{+}\boldsymbol{pAKT}^{\boldsymbol{na}\boldsymbol{15}}}}^{\boldsymbol{na}\boldsymbol{15}}$ | |
| associated reverse reaction | $\boldsymbol{v}\boldsymbol{16=k}\boldsymbol{16*}\frac{\boldsymbol{pp}\boldsymbol{53}}{\boldsymbol{pp}\boldsymbol{53+Km}\boldsymbol{16}}$ |  |
| p53 synthesis | $\boldsymbol{v}\boldsymbol{17=k}\boldsymbol{17}$ |  |
| MDM2 synthesis | $\boldsymbol{v}\boldsymbol{18=k}\boldsymbol{18*}\frac{{\boldsymbol{(p}\boldsymbol{53+pp}\boldsymbol{53)}}^{\boldsymbol{2}}}{{\boldsymbol{(p}\boldsymbol{53+pp}\boldsymbol{53)}}^{\boldsymbol{2}}\boldsymbol{+Km}\boldsymbol{18}}$ | |
| p53 degradation | $\boldsymbol{v}\boldsymbol{19=k}\boldsymbol{19*}\frac{\boldsymbol{p}\boldsymbol{53}}{\boldsymbol{1+}\frac{{\boldsymbol{Km}\boldsymbol{19}}^{\boldsymbol{2}}}{{\boldsymbol{MDM}\boldsymbol{2}}^{\boldsymbol{2}}}}$ |  |
| pp53 degradation | $\boldsymbol{v}\boldsymbol{20=deg}\boldsymbol{2*k}\boldsymbol{19*pp}\boldsymbol{53}$ |  |
| MDM2 degradation | $\boldsymbol{v}\boldsymbol{21=k}\boldsymbol{21*MDM}\boldsymbol{2}$ |  |
| iII) Algebraic Constraints |  |  |
| Active p38 | $\boldsymbol{ap}\boldsymbol{38=SP*pp}\boldsymbol{38}$ |  |
| Unphosphorylated p53 | $\boldsymbol{p}\boldsymbol{53}\boldsymbol{=p}\boldsymbol{53}\boldsymbol{t-pp}\boldsymbol{53}\boldsymbol{,}$ | where $\boldsymbol{p}\boldsymbol{53}\boldsymbol{t}$ denotes the total p53 protein concentration |
| The parameters k17 and k18 were calculated as a function of the p53 and MDM2 initial conditions ($\bar{\boldsymbol{p}\boldsymbol{53}}\boldsymbol{,}\bar{\boldsymbol{MDM}\boldsymbol{2}}$) assuming steady state for $\boldsymbol{doxo=0}$, see *Methods*. | | |
| Rate constant of p53 synthesis | $\boldsymbol{k}\boldsymbol{17= k}\boldsymbol{19}\bar{\boldsymbol{p}\boldsymbol{53}}\frac{\boldsymbol{Km}\boldsymbol{19}^{\boldsymbol{2}}}{\boldsymbol{Km}\boldsymbol{19}^{\boldsymbol{2}}\boldsymbol{+}{\bar{\boldsymbol{MDM}\boldsymbol{2}}}^{\boldsymbol{2}}}$ | |
| Rate constant of MDM2 synthesis | $\boldsymbol{k18=}\bar{\boldsymbol{MDM}\boldsymbol{2}}\boldsymbol{*k}\boldsymbol{21*}\frac{{\bar{\boldsymbol{p}\boldsymbol{53}}}^{\boldsymbol{2}}\boldsymbol{+Km}\boldsymbol{18}}{{\bar{\boldsymbol{p}\boldsymbol{53}}}^{\boldsymbol{2}}}$ | |
| The parameters k8 and k14 were calculated as a function of the AKT,PI3K,pAKT and pPI3K initial conditions ($\bar{\boldsymbol{AKT}}\boldsymbol{0,}\bar{\boldsymbol{PI}\boldsymbol{3}\boldsymbol{K}\boldsymbol{0}}\boldsymbol{,}\bar{\boldsymbol{pAKT}}\boldsymbol{0,}\bar{\boldsymbol{pPI}\boldsymbol{3}\boldsymbol{K}\boldsymbol{0}}$) assuming steady state for $t=0$ and therewith that $\boldsymbol{v}\boldsymbol{7=v}\boldsymbol{8}$ and $\boldsymbol{v}\boldsymbol{13=v}\boldsymbol{14}$, see *Methods*. | | |
| Rate constant of PI3K dephosphorylation | $\boldsymbol{k}\boldsymbol{8 =}\frac{\left( \bar{\boldsymbol{pPI}\boldsymbol{3}\boldsymbol{K}\boldsymbol{0}}\boldsymbol{+Km}\boldsymbol{8} \right)\boldsymbol{*k}\boldsymbol{7*}\bar{\boldsymbol{PI}\boldsymbol{3}\boldsymbol{K}\boldsymbol{0}}}{\left( \bar{\boldsymbol{PI}\boldsymbol{3}\boldsymbol{K}\boldsymbol{0}}\boldsymbol{+Km}\boldsymbol{7} \right)\boldsymbol{*}\bar{\boldsymbol{pPI}\boldsymbol{3}\boldsymbol{K}\boldsymbol{0}}}$ | |
| Rate constant of AKT dephosphorylation | $\boldsymbol{k}\boldsymbol{14 =}\frac{\boldsymbol{LY*k}\boldsymbol{13*}\bar{\boldsymbol{pPI}\boldsymbol{3}\boldsymbol{K}\boldsymbol{0}}\boldsymbol{*}\bar{\boldsymbol{AKT}\boldsymbol{0}}\boldsymbol{* (}\bar{\boldsymbol{pAKT}\boldsymbol{0}}\boldsymbol{+Km}\boldsymbol{14)}}{\boldsymbol{(}\bar{\boldsymbol{AKT}\boldsymbol{0}}\boldsymbol{+Km}\boldsymbol{13)*}\bar{\boldsymbol{pAKT}\boldsymbol{0}}}$ | |
| Differential equations |  | |
| $\frac{\boldsymbol{d}}{\boldsymbol{dt}}\boldsymbol{ATM}\boldsymbol{=}$ | $\boldsymbol{v}\boldsymbol{2-v1}$ | |
| $\frac{\boldsymbol{d}}{\boldsymbol{dt}}\boldsymbol{p}\boldsymbol{ATM}\boldsymbol{=}$ | $\boldsymbol{v1-v2}$ | |
| $\frac{\boldsymbol{d}}{\boldsymbol{dt}}\boldsymbol{MKK}\boldsymbol{4/7} \boldsymbol{=}$ | $\boldsymbol{v4-v3}$ | |
| $\frac{\boldsymbol{d}}{\boldsymbol{dt}}\boldsymbol{pMKK}\boldsymbol{4/7} \boldsymbol{=}$ | $\boldsymbol{v3-v4}$ | |
| $\frac{\boldsymbol{d}}{\boldsymbol{dt}}\boldsymbol{JNK} \boldsymbol{=}$ | $\boldsymbol{v10-v9}$ | |
| $\frac{\boldsymbol{d}}{\boldsymbol{dt}}\boldsymbol{pJNK} \boldsymbol{=}$ | $\boldsymbol{v9-v10}$ | |
| $\frac{\boldsymbol{d}}{\boldsymbol{dt}}\boldsymbol{MKK}\boldsymbol{3/6} \boldsymbol{=}$ | $\boldsymbol{v6-v5}$ | |
| $\frac{\boldsymbol{d}}{\boldsymbol{dt}}\boldsymbol{pMKK}\boldsymbol{3/6} \boldsymbol{=}$ | $\boldsymbol{v5-v6}$ | |
| $\frac{\boldsymbol{d}}{\boldsymbol{dt}}\boldsymbol{p}\boldsymbol{38} \boldsymbol{=}$ | $\boldsymbol{v1}\boldsymbol{2}\boldsymbol{-v1}\boldsymbol{1}$ | |
| $\frac{\boldsymbol{d}}{\boldsymbol{dt}}\boldsymbol{pp}\boldsymbol{38} \boldsymbol{=}$ | $\boldsymbol{v}\boldsymbol{1}\boldsymbol{2-v1}\boldsymbol{2}$ | |
| $\frac{\boldsymbol{d}}{\boldsymbol{dt}}\boldsymbol{PI}\boldsymbol{3}\boldsymbol{K} \boldsymbol{=}$ | $\boldsymbol{v}\boldsymbol{8}\boldsymbol{-v}\boldsymbol{7}$ | |
| $\frac{\boldsymbol{d}}{\boldsymbol{dt}}\boldsymbol{pPI}\boldsymbol{3}\boldsymbol{K} \boldsymbol{=}$ | $\boldsymbol{v}\boldsymbol{7}\boldsymbol{-v}\boldsymbol{8}$ | |
| $\frac{\boldsymbol{d}}{\boldsymbol{dt}}\boldsymbol{A}\boldsymbol{KT} \boldsymbol{=}$ | $\boldsymbol{v}\boldsymbol{14}\boldsymbol{-v1}\boldsymbol{3}$ | |
| $\frac{\boldsymbol{d}}{\boldsymbol{dt}}\boldsymbol{pAKT} \boldsymbol{=}$ | $\boldsymbol{v}\boldsymbol{13}\boldsymbol{-v}\boldsymbol{14}$ | |
| $\frac{\boldsymbol{d}}{\boldsymbol{dt}}\boldsymbol{p}\boldsymbol{53}\boldsymbol{t} \boldsymbol{=}$ | $\boldsymbol{v}\boldsymbol{17}\boldsymbol{-v1}\boldsymbol{9-v}\boldsymbol{20}$ | |
| $\frac{\boldsymbol{d}}{\boldsymbol{dt}}\boldsymbol{pp}\boldsymbol{53} \boldsymbol{=}$ | $\boldsymbol{v}\boldsymbol{15}\boldsymbol{-v1}\boldsymbol{6-v}\boldsymbol{20}$ | |
| $\frac{\boldsymbol{d}}{\boldsymbol{dt}}\boldsymbol{MDM}\boldsymbol{2} \boldsymbol{=}$ | $\boldsymbol{v}\boldsymbol{18}\boldsymbol{-v}\boldsymbol{21}$ | |
